# Mobile genetic elements that shape microbial diversity and functions in thawing permafrost soils

**DOI:** 10.1101/2025.02.12.637893

**Authors:** Jiarong Guo, Samuel Aroney, Guillermo Dominguez-Huerta, Derek Smith, Dean Vik, Carlos Owusu Ansah, Akbar Adjie Pratama, Sergei Solonenko, Funing Tian, Cristina Howard-Varona, ZhiPing Zhong, Aminata Fofana, Garrett Smith, Suzanne B. Hodgkins, Dylan Cronin, EMERGE Field Teams 2010-2019, EMERGE Coordinators, Ben J. Woodcroft, Gene W. Tyson, Virginia I. Rich, Matthew B. Sullivan, Simon Roux, Sarah C. Bagby

## Abstract

The world’s ecosystems are shaped by microbiota. Their niches and their impacts depend on functional profiles influenced by gene gains and losses. While culture-based experiments demonstrate that mobile genetic elements (MGEs) can mediate gene flux, quantitative field data on the rates and impacts of MGE activity remains scarce. Here we leverage large-scale soil meta-omic data to develop and apply analytics for studying MGEs in complex natural systems. In our model permafrost-thaw ecosystem, Stordalen Mire, we identify ∼2.1 million MGE recombinases across 89 microbial phyla to assess ecological distributions, affected functions, past mobility, and current activity. This revealed MGEs shaping natural genetic diversity via differential impacts on major phyla; affecting a wide range of functions, including diverse regulatory and metabolic genes affecting carbon flux and nutrient cycling; and moving at rates that should significantly influence the realized functional profiles of natural microbiomes. These findings and this systematic meta-omic framework open new avenues to better investigate MGE diversity, activity, mobility, and impacts in nature.

## Main text

Microbiomes are increasingly recognized for driving biogeochemical and nutrient cycling across diverse systems (*1–4*). As metagenomics has advanced, our understanding of microbial diversity in these systems has sharpened. Microbial diversity was first explored via low-resolution methods like denaturing gradient gel electrophoresis and 16S amplicon sequencing (reviewed in (*5*)), then came into focus through shotgun metagenomics that enabled the reconstruction of entire metagenome-assembled genomes (MAGs) from these uncultivated microbes (*6*). To obtain these genomes, metagenome binning approaches typically collapse diversity within a given population, with variant sequences omitted from the binned consensus (*7*). Emerging tools can profile strain-level diversity in metagenomic datasets from intensively studied systems like the human microbiome, but still struggle in complex communities (e.g., soils) and those with limited reference sequences (*8–10*). Yet within-species variation can have profound effects on ecosystem outcomes, as in well-documented cases of commensal and pathogenic gut microbiota separated only by strain-level variation (e.g., (*11*)). Such microdiversity can arise both through mutation and through gene flow ((*12*); reviewed in (*13*)). Understanding the constraints on these processes is a critical step towards understanding the pace and scope of adaptive change accessible to microbiomes in changing environments.

Mobile genetic elements (MGEs) are a major driver of gene flow and genomic rearrangement (*14, 15*). Recombinase genes enable MGEs to move between genomic loci and can be used to distinguish between several MGE types (*16*): insertion sequences and transposons (“IS_Tn”), phage and phage-like elements (“Phage”), conjugative elements and mobility islands (“CE”), and integrons. These types differ in several respects, including their ability to move between cells unaided and the likelihood that, when they move, they carry genes that are not required for MGE mobility but that may alter host metabolism or function when expressed (“cargo genes”). MGEs can influence host phenotypes and niches through their cargo and/or the host genomic context of MGE insertion (Fig. 1A), causing gains or losses of function and/or regulatory shifts (*17, 18*) that should scale and diversify with the number of such movements (Fig. 1B). The wider an MGE’s host range is, the greater is its range of partners—both hosts and other MGEs—for exchanging genetic modules (*19*).

**Figure 1.**
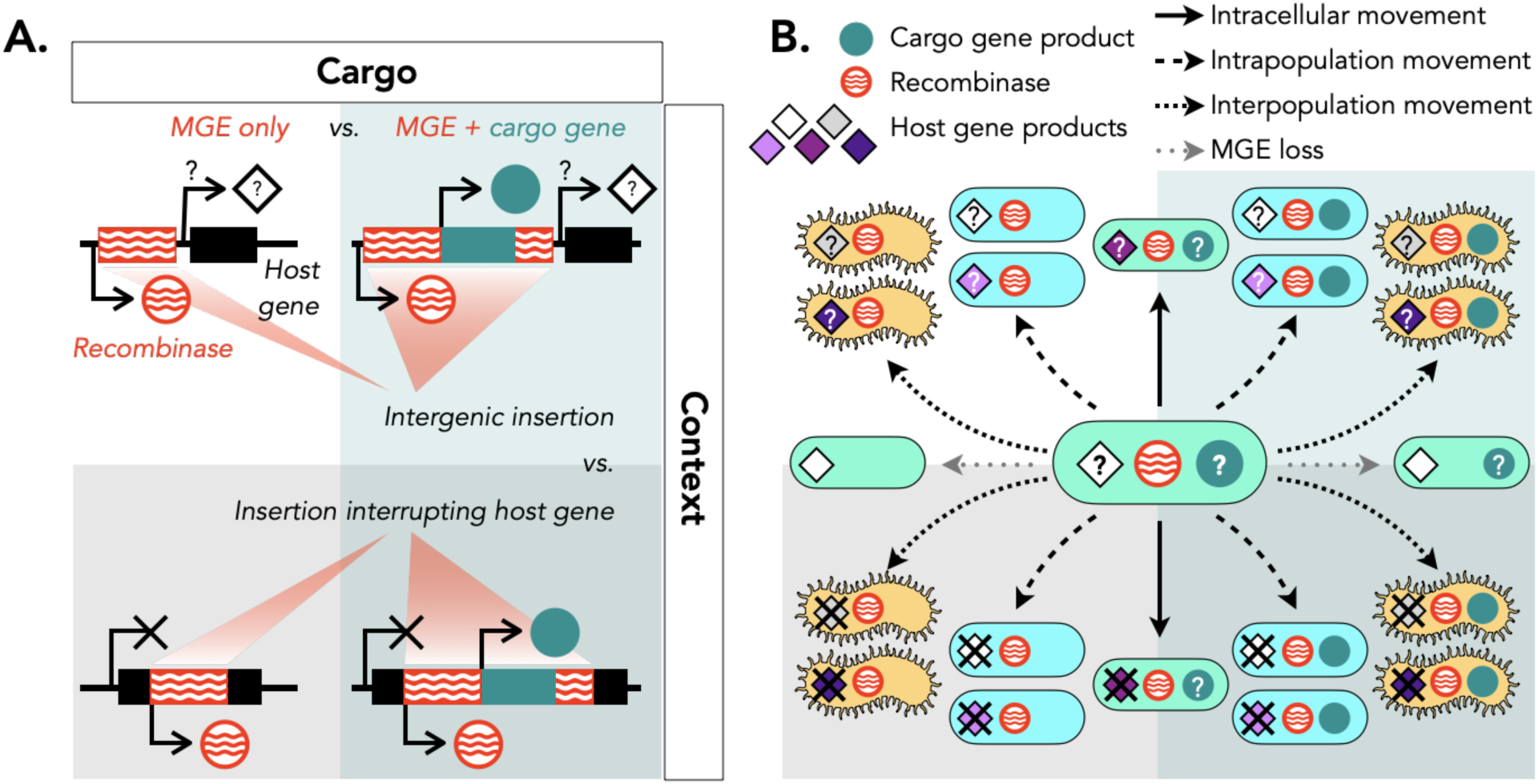
MGE cargo and context determine the impact of MGE activity on host coding capacity and population genetic diversity. **A.** An MGE may or may not carry cargo and may insert either intergenically, potentially altering transcriptional regulation downstream, or into a host gene, causing function loss. MGE and recombinase gene product, red and white waves; cargo gene and product, solid teal; host gene, black; host gene product, white diamond. Disrupted genes yield no product (X); genes downstream of MGE insertions may or may not be transcribed (?). **B.** Each cycle of excision and integration may shift the MGE and its insertion site and/or cargo between contexts, generating new genetic diversity. Intracellular movement (solid arrows, teal cells) may interrupt (X) or influence gene regulation of (?) a new host gene (diamonds); intrapopulation movement (dashed arrows, blue cells) lets the MGE interrupt or influence the same or different host genes in a new genetic background; interpopulation movement (dot-dashed arrows, yellow cells) lets the MGE interrupt or influence a new suite of genes; and MGE loss (gray dotted arrows) may remove or leave behind cargo genes and potentially restore function of previously disrupted host genes. Not depicted: capture of host genes as new cargo genes.

Yet to date these impacts have not been accessible to comprehensive characterization in a natural, non-host-associated community. With comparatively few reference genomes available in such communities, analysis depends on de novo assembly, but the mobility of MGEs tends to break assembly graphs (*20, 21*); where binning succeeds, the loss of strain-level variation in consensus sequences hides differences in MGE presence, absence, or genomic context. Moreover, few projects incorporate the long-term repeated deep sequencing that could reveal MGE activity over time. These structural challenges have made MGEs an elephant in the room, a complication that natural-system metagenomics has largely had to accept as an unknown. Targeted studies have grappled with some aspects of MGE biology: for instance, numerous studies have illuminated transfers of a given MGE-associated trait (e.g., antimicrobial resistance (*16, 22, 23*)) or a given MGE type (*24–26*), and substantial progress has been made in identifying MGEs of different types in the genomes of isolates (*16*). But to date there is no comprehensive survey of MGEs in a complex natural system, and thus no quantitative basis for bounding their potential impacts. The minimal metagenomic dataset that can support such analyses must deeply sequence the same microbiome repeatedly over short and long timescales.

Long-term ecological study sites provide an opportunity to capture such datasets. The model permafrost peatland Stordalen Mire, Sweden, lies in the rapidly changing discontinuous permafrost margin, where rising temperatures convert the intact palsa habitat to partially thawed bog and then fully thawed fen, with concomitant increases in methane efflux from the soil (**Fig. 2A**) (*27–29*). Soil sampling in these three habitats over almost a decade of peak growing seasons (2010–2017, 2019) has yielded 615 short-read metagenomes (*30*), of which 50 have paired short-read metatranscriptomes and 6 have paired long-read metagenomes. Previous analyses of a subset of this data have shed light on key taxa and processes in the system (*28*) and have pointed to the role of functional redundancy and dispersal in allowing each habitat’s microbiome to resist environmental change (*28, 31*). MGE-mediated gene flow could have synergistic effects with both, but to date the analytic approaches needed even to bound the potential impacts of MGEs on a natural system have been lacking.

**Figure 2.**
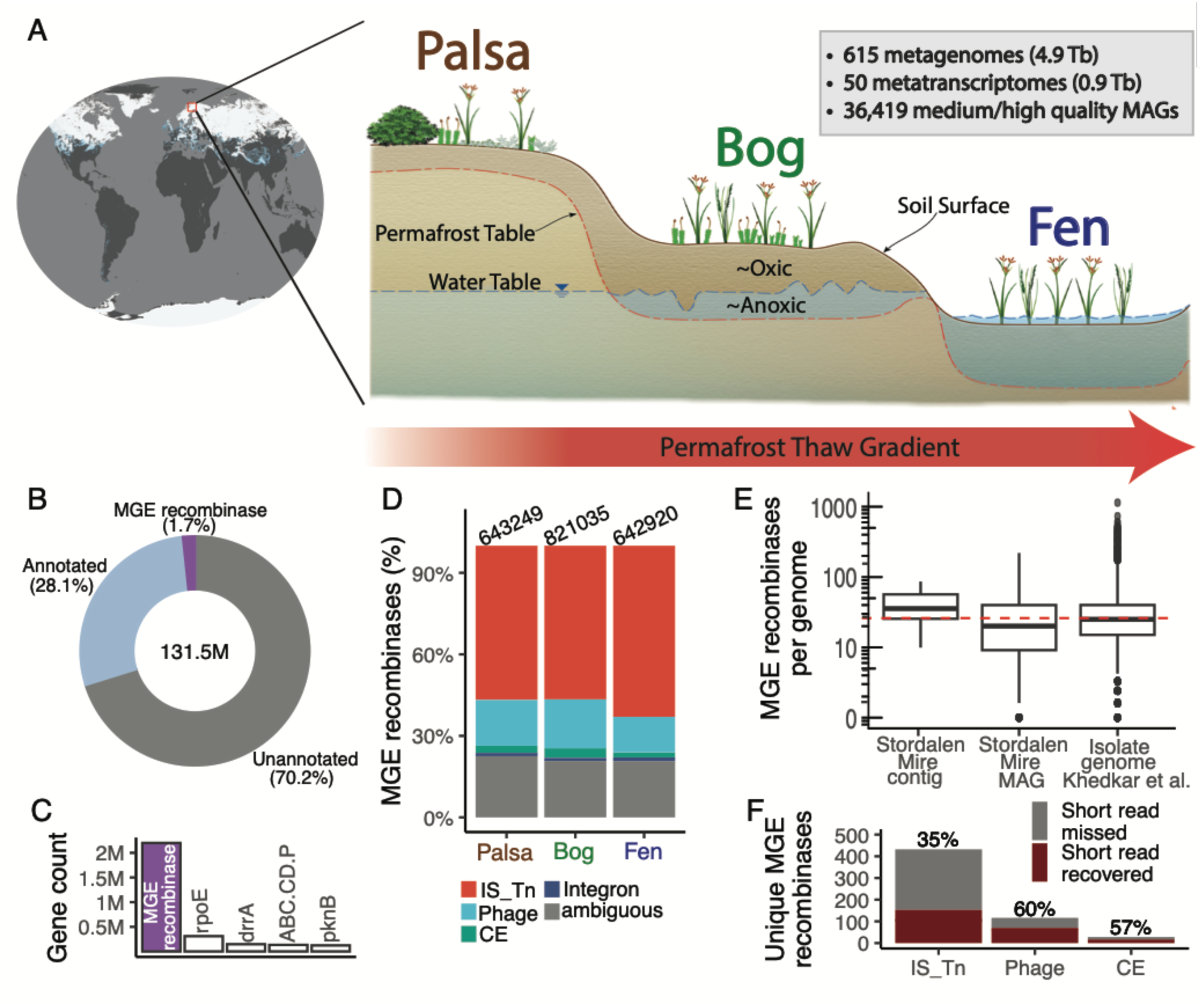
The Stordalen Mire model ecosystem and its mobile genetic element (MGE) recombinases. **A.** Long-term study site (image credit: NASA) with large datasets (gray inset) along a palsa-bog-fen thaw gradient. **B.** MGE recombinase percentage in all genes from Stordalen contigs (≥1,500 bp). **C**. Counts of most abundant genes: MGE recombinase; *rpoE*, RNA polymerase sigma 70 factor; *drrA*, antibiotic transport system ATP binding protein; *pknB*, serine/threonine protein kinase; *ABC.CD.P*, putative ABC transport system permease. **D.** MGE recombinase type distributions: “IS_Tn”, insertion sequence/transposon; “Phage”, phage and phage-like elements; “CE”, conjugative elements/mobility islands; “Integron”, integron; “ambiguous”, recombinases found in >1 MGE types. **E.** MGE estimates per genome for Stordalen Mire contigs (median=36) or MAGs (median=23) and from cultured isolates in Khedkar et al. (median=25; red dashed line). Filled circles, points >1.5 times the interquartile range below the first quartile or above the third quartile. **F.** Comparison of MGE recombinase recovery between short-read and hybrid assemblies (n = 17), with recombinase matching at 100% protein identity. “Short read missed”: recombinase found only in hybrid MAGs; “short read recovered”: recombinase found in both hybrid and short-read MAGs (recovery rates shown as percentages on top of bars).

Here we use the rich Stordalen Mire dataset to develop and apply these metrics. We use paired short- and long-read sequencing to determine the expected yields in MGE recovery from short-read data, and use these rates to obtain conservative bounds on the per-genome abundance of MGEs of different types across different phyla. We assess the functions of genes stringently identified as affected by MGEs, revealing that MGE-mediated gene gain and loss reaches many functions in core nutrient metabolism with implications for community metabolism and ecosystem outputs. Finally, we present three complementary metatranscriptomic and metagenomic metrics of MGE mobility at different timescales and across MGE types. Collectively, these analyses shed light on the scale and potential impacts of MGE activity in a soil microbiome and provide a set of approaches that will enable quantitative investigation of MGEs in short-read metagenomic data from complex natural systems.

### MGE recombinases from permafrost thaw gradient microbiomes

Stordalen Mire is a climate-critical permafrost peatland, extensively studied for decades to assess the interplay of vegetation, microbiota, and climate along a three-stage thaw gradient of palsa, bog, and fen habitats that represent, respectively, the least to most thawed soils. Prior to our work, ∼5.8 terabases of metagenomic and metatranscriptomic sequencing data had been generated from 8 years of field sampling (**Fig. 2A, table S1**). We first identified and quantified all major MGEs in these data via recently developed hidden Markov models (HMMs) and MGE identification protocols that leverage recombinases as MGE hallmark genes (*32*) (**table S2**). Specifically, we screened 131.5M genes predicted from 30.7M short-read metagenomic contigs, and this identified 2,107,204 MGE recombinases (1,130,776 unique protein sequences) derived from 1,790,672 contigs (**Fig. 2B,C&D, Extended Data fig. 1, Extended Data fig. 2A&B, Extended Data fig. 3A,B, table S3 item A**). On average, this represented 17.3 MGE recombinases per million base pairs (Mbp) of microbiome sequence data, dwarfing the representation found for even the most abundant KEGG gene family, the stress-response sigma factor *rpoE* (2.4 copies/Mbp; **Fig. 2C**).

We next classified these abundant recombinases into four previously-established types (*32*): (i) integrons, (ii) insertion sequences and transposons, herein “IS_Tn”, (iii) phage / phage-like elements, herein “Phage”, and (iv) conjugative elements and mobility islands, herein “CE”. Classification facilitates evaluation of ecological impacts, since these types are known to differ in length, mode of propagation, and gene content. Prior work evaluating 187 metagenomes available in 2010 and >80K bacterial and archaeal isolate genomes (*32, 33*) found IS_Tn dominant in highly abundant MGE pools. In Stordalen Mire, we also found thatIS_Tn was the most commonly recovered type (58.6%), distantly followed by Phage (16.1%), CE (2.8%), and integron (1.2%) (the remaining 21.3% were ambiguous; see Methods; **Extended Data fig. 2A**). Ecologically, at this high level of recombinase types, this rank-order pattern remained remarkably constant across time (8 years of peak growing season sampling) and space (within and between the study site’s three habitat types) (**Fig. 2D, Extended Data fig. 2B,C**). The same pattern largely persisted across phyla for both MAG- and contig-level taxonomic assignments (216,720 and 804,456 MGE recombinases, respectively, assigned to a total of 89 phyla with 93.7% agreement in assignments for the subset shared; **Fig. 3A,B**, **Extended Data fig. 4A,B, table S3 item A**). MGE distribution broadly followed host lineage relative abundances, with most MGEs in ecosystem-dominant bacterial and archaeal phyla such as Acidobacteriota, Actinomycetota, Pseudomonadota, Verrucomicrobiota, and Halobacteriota (*28*) (**Fig. 3**, **Extended Data figs. 4, 5**). Echoing isolate-based observations (*32*), the virtual absence of integrons in the abundant Actinomycetota stands out against a background of low cross-phylum variation in the frequency of each MGE type in our Stordalen microbiomes.

**Figure 3.**
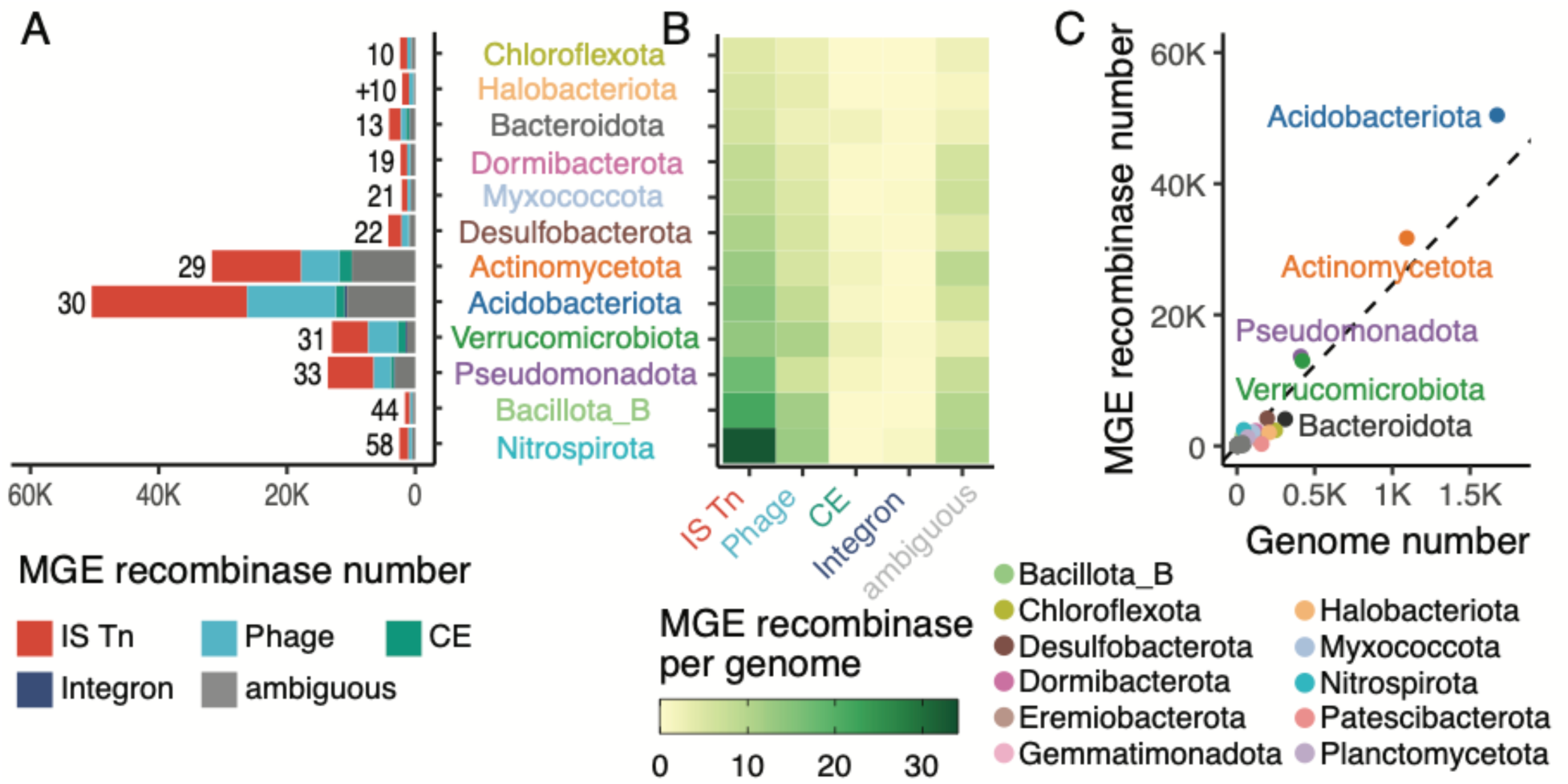
MGE recombinase type distribution across microbial host lineages (MAG-based) of the full Stordalen Mire short read dataset. **A.** Number of each MGE type found in each phylum, as identified from contigs binned into MAGs. MGE recombinase counts shown for the top 12 phyla (out of 46 total), ordered by average MGE recombinase number per genome (indicated to the left of each bar with “+” indicating Archaea). **B.** Number of MGE recombinases identified per genome grouped by MGE type and host phylum. **C.** Number of MGE recombinases (y-axis) against estimated number of genomes per phylum (x-axis). Dotted line represents the global phylum-averaged number of MGE recombinases per genome.

Next, we wondered how the MGE frequency of ∼25 MGEs/genome measured in isolates would compare with estimates from natural populations, given the potential for MGEs to be lost during laboratory cultivation due to reduced selective pressure in clonal isolates. To this end, we quantified MGEs per genome via two approaches applied to our Stordalen microbiomes. First, we used a contig-based estimate, comparing identified MGE recombinase counts to total genome counts estimated using a standard single-copy marker gene approach for each sample (*34, 35*). This revealed a median of 36 MGE recombinases per genome in Stordalen microbiomes (**Fig. 2E**). Separately, we considered only the subset of MGEs that were binned into medium/high-quality metagenome-assembled genomes (MAGs) (≥70% completeness, ≤10% contamination) (**table S3 item B**). Scaling per-MAG MGE counts by MAG completeness revealed a median of 23 MGE recombinases per genome (**Fig. 2E**). Comparing the two approaches, we note that only 15.9% of MGE-bearing contigs were binned, even though Stordalen MAGs capture a remarkable 73% of sequence reads at genus level (*28, 31*). Thus, we interpret the MAG-based estimate as a lower bound since it empirically captures only a subset of the contig-observed MGEs, and dynamic rearrangements in MGE-bearing genomic regions are a known challenge for short-read metagenome assembly and binning (*36, 21, 37*).

To empirically quantify the impact of assembly and binning challenges on short-read counts, we performed long-read sequencing on six samples and evaluated MGE recombinase recovery across the assembly and binning process. In total, these data suggested that short-read assembly and binning recovered 35–60% of MGE recombinases, with IS_Tn recombinases disproportionately under-recovered (**Fig. 2F**, **Extended Data figs. 6, 7; Supplementary text**). Using this benchmarking to scale our MAG-based counts, we obtain a revised estimate of 38–66 MGE recombinases per genome *in situ*. We interpret this number as comparable with the contig-based estimate and consistent with the hypothesis that isolates might lose MGEs during lab cultivation.

### MGE impacts range far beyond biotic social interactions

We next wondered whether MGEs affected genes of ecological importance, and how these fit into microbiome activity in Stordalen Mire. Prior genome-resolved metagenomic analyses (*28, 31*) show that intercellular interactions and public goods matter in Stordalen Mire: cross-feeding relationships between carbon generalists and methanogens are persistent and strongly associated with methane flux, and extracellular degradation of polysaccharides is a bottleneck in carbon flow. Beyond microbes, there are also abundant viral community members that harbor auxiliary metabolic genes that likely mediate exopolysaccharide breakdown (*29*). Such outward-directed “social” functions are expected to be affected by MGEs (*38–41*), as are some stress responses (*42*), but these are only a small slice of microbial activities. Each cell must also manage its own routine “domestic” processes (e.g., genome maintenance, gene expression, nutrient homeostasis) (*43–46*), functions that shape physiology and ecosystem outputs but that have been little studied in the mobilome to date. Recently observed plasmid-borne genes affecting domestic activities like nitrogen fixation (*42*) highlight the potential for MGEs to influence nutrient cycling directly, but the prevalence of such functions among MGE-affected genes remains unknown.

Thus, we asked which Stordalen Mire functions could be confidently detected as mobilome-affected (i.e., encoded in MGEs or interrupted by them) and how far beyond biotic interactions the mobilome’s direct influence extends. To ensure that host gene activities were not misattributed to MGEs, we adopted a highly conservative approach for curating MGE boundaries. It should be noted that only ∼0.8% of the total 2.1M MGE recombinases identified were eligible for analysis of MGE-affected functional genes under this stringent approach. Annotation identified putative functions for cargo genes within conservatively defined boundaries of 5,288 MGEs (900 CEs, 551 integrons, and 3,837 phage) and for host genes interrupted by 11,295 IS_Tns. (**Extended Data fig. 1, Methods**). Among this set of MGEs, our manually curated annotation identified high-confidence functional predictions for 3,631 MGE-affected genes with 1,199 distinct KEGG annotations (**table S3 item C-F**, **table S4, Supplementary text**).

As expected, this reduced set of curated MGE-affected genes included many related to host biotic interactions (n=1377; **Fig. 4, table S4**). This included diverse phage defense systems (379 carried, 168 interrupted) that can abort infection, degrade phage DNA by restriction-modification, or inhibit phage DNA synthesis by nucleotide depletion (dCTP deaminase) (*40, 47*). MGEs also impact genes for YD-repeat-containing proteins (10 carried, 29 interrupted), a contact-dependent growth inhibitor thought to shape responses to high population densities (*48*). In social interactions mediated by public goods, MGEs impact genes encoding carbohydrate-active enzymes (CAZymes) involved in synthesis or degradation of extracellular polysaccharides (154 carried, 125 interrupted). The former produce public goods and molecules involved in cell-phage and cell-cell interactions, while the latter can become public goods through direct secretion (extracellular enzymes) or outer membrane vesicle budding (periplasmic enzymes) (*49*). Separately, MGEs appear to impact synthesis and transport of other public goods (39 carried, 38 interrupted), predominantly siderophores. Finally, MGEs can impact genes that encode a public service (2 carried, 13 interrupted), the detoxification of diffusible peroxides (*50*).

**Figure 4.**
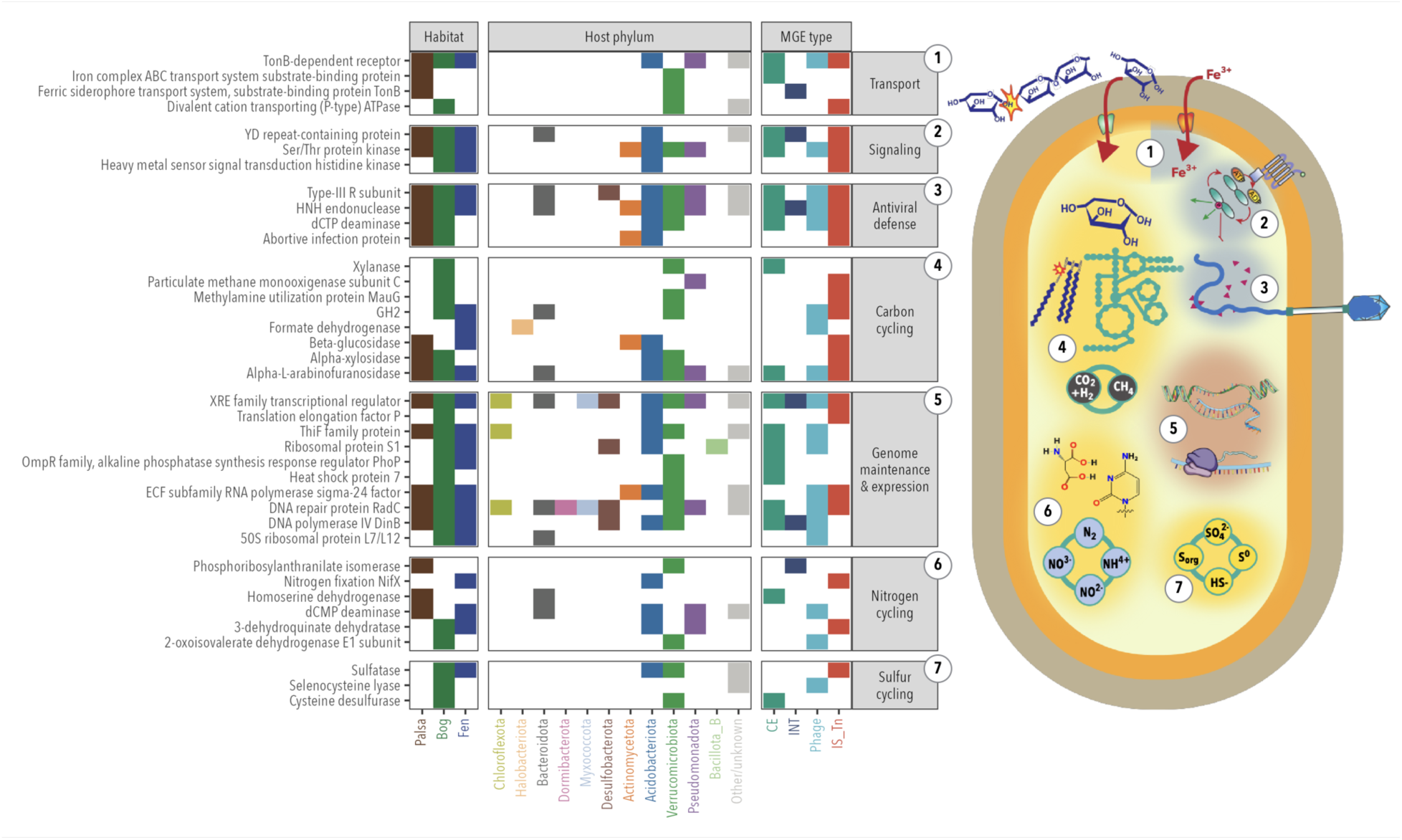
Host functions impacted by MGEs in Stordalen Mire. Distribution across habitats, host phyla, and MGE types of selected nutrient cycling pathways (yellow highlights in cell, groups 1, 4, 6, and 7), housekeeping functions (red highlight, group 5), and signaling and social functions (blue highlights, groups 1–3) impacted by MGEs whose boundaries could be conservatively established. Colored boxes indicate positive detection of an MGE-affected function in a habitat, phylum, or MGE type. Phyla are ordered from left to right by increasing per-genome MGE count (see fig. 3). CE, INT, and phage carry impacted functions as cargo genes; IS_Tn interrupt host genes, causing loss of function.

Beyond biotic interactions, we identified diverse MGEs seemingly affecting domestic processes. For example, we uncovered MGEs affecting central carbon metabolism genes (6 carried, 44 interrupted; **table S4**). Intriguingly, even in this small subset of conservatively-curated MGEs, there were three direct connections to methane metabolism: the key hydrogenotrophic metabolism gene formate dehydrogenase (*51*) was carried by a Phage from Halobacteriota (*Methanoregula* sp003136075), a group of methanogens active at Stordalen Mire (*27, 28, 31*), and IS_Tns interrupted a methane monooxygenase (*pmoC*) subunit in a Pseudomonadota (unbinned Alphaproteobacterial contig) and the methylamine utilization gene *mauG* in a Verrucomicrobiota (unbinned UBA11358 contig). Thus, even within this highly conservative subset of MGE-affected genes, we find genomic evidence that MGEs may alter Stordalen methane fluxes both directly (by changing genomic potential for methanogenesis and methanotrophy) and indirectly (by changing the distribution of carbon-processing pathways that control the flow of substrates to methanogens).

MGEs also might influence other major nutrient cycles, trace metal cycles, and cellular housekeeping functions. In nitrogen cycling, we found MGEs influencing nitrogen processing (26 carried, 58 interrupted; **Fig. 4, table S4**) across the thaw gradient, predominantly in amino acid metabolism but also touching other nitrogen processes, including fixation. Because fixed nitrogen is thought to be a partially “leaky” public good (*52*), IS_Tn interruption of an Acidobacterial nitrogen fixation regulator (*53*) illustrates the potential for MGEs influencing growth beyond the host cell by controlling nitrogen. The CAZyme genes highlighted above include 10 chitinase genes carried and 1 interrupted by MGEs; chitin is hypothesized to serve as a key alternative nitrogen source at our N-limited site (*31*). In sulfur cycling, we found MGEs that affect genes putatively related to inorganic sulfur limitation (19 carried, 15 interrupted), including sulfatases that might degrade sulfated polysaccharides (*44*) and cysteine desulfurases and selenocysteine lyases that might scavenge thiol groups and reduce sulfur demand (*54, 55*). Beyond macronutrient cycling, MGEs impact iron cycling (39 carried, 37 interrupted) via siderophore production and transport, roles highlighted above in connection with public goods. Finally, MGEs also encode core housekeeping genes involved in genome maintenance: DNA repair, predominantly *radC* (89 carried, 49 interrupted; genome replication, predominantly *dinB* and *polC* (79 carried, 15 interrupted); regulation of host gene transcription (RNA polymerase subunits, transcriptional regulators; 174 carried, 135 interrupted), and translation (ribosomal proteins and one elongation factor; 32 carried, 14 interrupted); and proteome maintenance (predominantly chaperonin, ubiquitin activation, and proteases; 104 carried, 53 interrupted).

Critically, even though our stringent analysis criteria and the limitations of functional gene annotation in understudied environments together reduced the set of MGE-affected genes analyzed to 0.02% of the total MGEs identified, we still find abundant examples of mobilome-affected social and domestic functions beyond previous observations (*56*). Our analysis adds many further examples of MGE-impacted functions including those that influence carbon flow through this climate-critical ecosystem, both directly (exopolysaccharide degradation, central carbon metabolism) and indirectly (via nitrogen, sulfur, and/or iron availability to support carbon flux). Although research to date has focused on MGE impacts on biotic interactions, MGE impacts on domestic functions appear relatively common, at the level of both the functions (∼51% domestic vs. ∼38% social; the remainder play both roles, support MGE maintenance and replication, or are uncategorized) and the MGEs themselves (∼55% affect at least one domestic function). If this conservative subset is representative of the Stordalen mobilome as a whole, then as sequencing methods, functional annotation, and databases improve, we would expect ∼1.1M of the full ∼2.1M MGE recombinases to influence domestic functions, with significant implications for nutrient cycling, as well as microbiome activity, ecology, and evolution.

### MGE recombinase mobility and activity

To assess ecological and evolutionary change in Stordalen Mire MGEs, we evaluated their mobility and activity from three different viewpoints: (i) snapshots of metatranscriptomic MGE recombinase expression as a proxy for mobilization underway, (ii) genomic neighborhood change as a marker of completed MGE movement between insertion sites, and (iii) partial carriage of an MGE by the host population as the result of evolution and selection over longer time scales. Integrating results across these analyses can help highlight individual MGE dynamics. For example, high recombinase expression rates together with low rates of genomic neighborhood change for a given MGE would suggest a strong preference among insertion sites. Our overall findings are as follows.

First, we asked whether MGE recombinases were “transcriptionally active” as measured via read mapping from 50 available metatranscriptomes against contigs from paired metagenomes (**table S5**), taking ribosomal gene mapping rates as a baseline for constitutive transcriptional activity of actively growing cells (*57*) (**table S6**). This revealed transcriptional activity in ∼3.4– 56.4% (median: 27.8%) of ribosomal genes and ∼0.04–9.1% (median: 1.2%) of MGE recombinase genes (**Fig. 5A, table S3 item G**), making MGE recombinases the second-most-transcribed non-ribosomal genes in Stordalen Mire (**Fig. 5A**). Stratified by recombinase type, we found roughly equal levels of transcriptional activity for IS_Tn and Phage recombinases versus ∼five-to ten-fold lower integron and CE recombinase transcription (**Fig. 5B**). Ecologically, IS_Tn recombinase transcription increased along the thaw gradient (significant from palsa to fen, p ≤ 0.01, Wilcoxon rank-sum test) (**Fig. 5C, fig.S8B**), suggesting increased MGE activity in the more-disturbed fen. Taxonomically, transcriptionally active MGE recombinases were detected across hosts spanning 30 phyla with activity largely proportional to MGE- and phylum abundance (**Fig. 5D, Extended Data fig. 8C**). However, one of these, the archaeal phylum *Halobacteriota,* was unique in that it is neither abundant nor a MGE recombinase-rich phylum, and yet was the most active phylum for MGE recombinase transcription. Given that Stordalen Mire *Halobacteriota* are known to be methanogens that are active and selected for in thawed fen and bog habitats (*27, 58*), we speculate that their disproportionate MGE recombinase activity might be adaptive.

**Figure 5.**
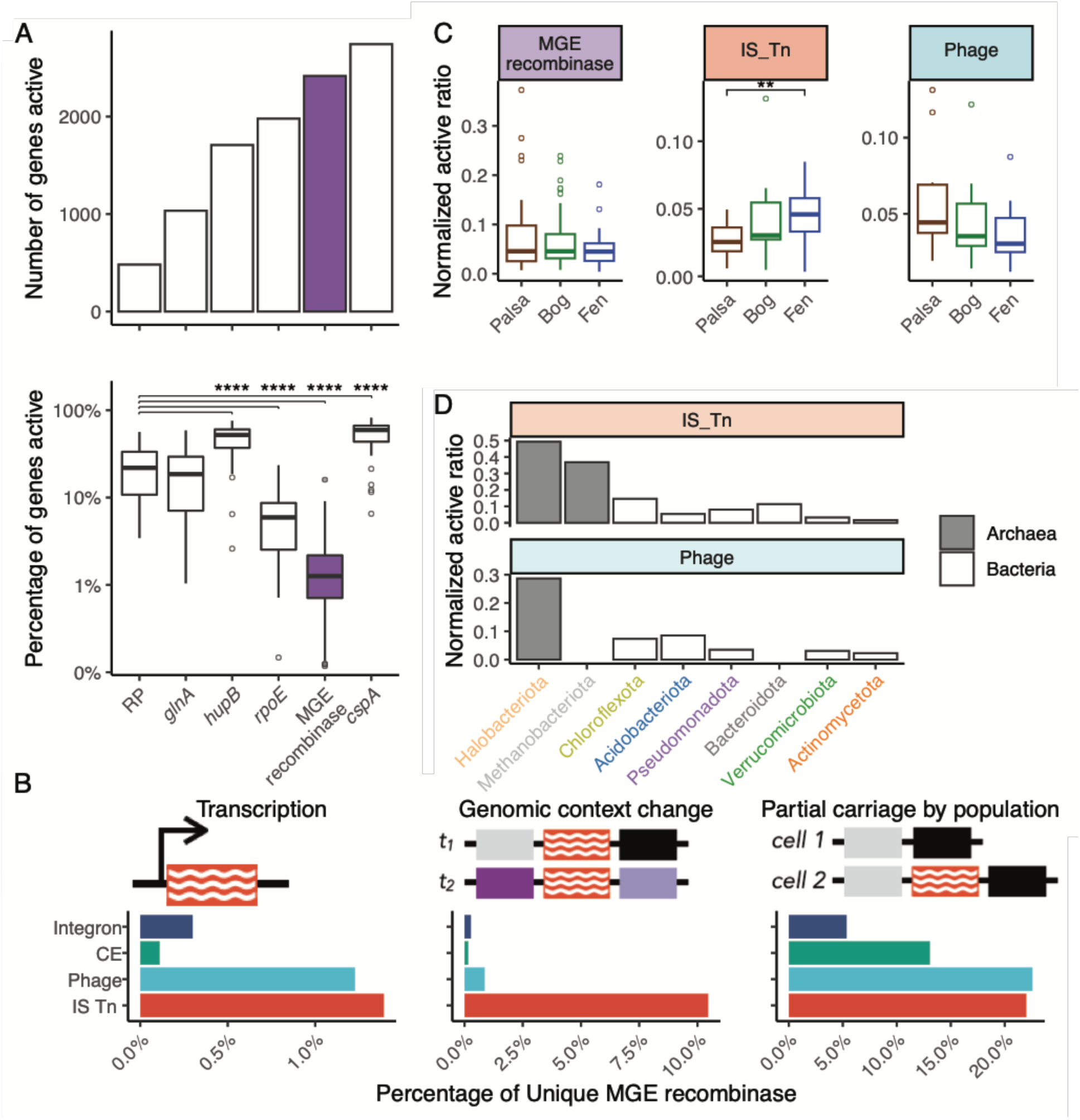
MGE recombinase activity. **A**. Top panel: Genes detected most frequently as active based on metatranscriptome read mapping. Gene categories from right to left: *cspA*, major cold shock protein gene; MGE recombinase; *rpoE*, RNA polymerase sigma-70 factor gene; *hupB*, DNA binding protein HU-beta gene; *glnA,* glutamine synthetase gene; RP, ribosomal protein genes. Bottom panel: percentage of genes active based on the full set of genes identified in the matching metagenome. Wilcoxon rank-sum test of statistical significance; ****, *P* value≤0.0001. **B**. Three facets of MGE recombinase activity/mobility--transcription (left), genomic neighborhood stability (middle), partial carriage by host population (right) – are conceptually presented (upper panel) and quantified and summarized per MGE recombinase type (lower panel). The x-axis is the percentage of unique MGE recombinase (100% identity at protein level) detected with activity / mobility within each type. **C**. Comparison of ratio of active MGE recombinase, “IS_Tn”, and “Phage” recombinases across three habitats, normalized by the ratio of active ribosomal genes for the same sample. **D.** Normalized active ratio of “IS_Tn” and “Phage” recombinase across host phyla with ≥5 active “IS_Tn” or “Phage” recombinases, and ribosomal genes. “CE” and “Integron” recombinases are not shown here due to low counts.

Second, where MGE recombinases were observed multiple times, we assessed the genomic context (300 bp upstream and downstream) surrounding each instance. We then interpreted this genomic context as a proxy for intra/inter-genomic mobility, either of the recombinase itself or of other recombinase-proximal genes (see examples of changing neighborhoods in **Extended Data fig. 9, Methods**). From the original ∼2.1M MGE recombinases, we identified 74,257 MGE recombinase clusters (100% average amino acid identity [AAI])) observed more than once in our dataset. Among these, most had identical genomic neighborhoods across observations, but some (0.1-10%, depending on MGE type) did show neighborhood variation (**Fig. 5B**, **table S3 item H**). Neighborhood variability was most common among IS_Tn recombinases (∼10%), and much less so for Phage (∼1%), integron (0.2%) and CE (0.1%) recombinases. This was consistent among years and habitats, with larger variation detected across diverse host phyla (up to 37% in Desulfobacterota_G) (**Extended Data fig. 10)**. Mindful that hypervariable islands in cellular genomes can often be impacted by multiple MGE types, we quantified co-occurring IS_Tn insertions, which could cause the neighborhood to change while the focal MGE remains stable. Fewer than 1 in 5 of our changed-genomic-neighborhood examples was due to co-occurring IS_Tn insertions (**Extended Data fig. 9**). Considering our approach only detects insertion in immediate neighborhoods of MGE recombinase, this is a lower bound estimate of mobility at hypervariable multiple-MGE genomic islands among MGE context changes at Stordalen.

Third, we sought to assess what fraction of a population contains any given MGE recombinase by evaluating read-mapping-based coverage of the recombinase against that of its surrounding genomic context (see Methods). Where MGE recombinase coverage was significantly lower, we infer that only a subset of the host population’s genotypes encoded the MGE recombinase, presumably a function of MGE mobility. Overall, across MGE types, we found 8,033 recombinase clusters (100% AAI) that had ≥20 kbp of genomic context. Of these, ∼5–23% showed variation in this genomic context consistent with at least part of the cellular population having the MGE and part not having the MGE (**Fig. 5B, table S3 item I**). These findings were again broadly consistent across years, habitats, and well-sampled lineages (**Extended Data fig. 11B-D**). In spite of spatio-temporal consistencies, there were notable variations in intrapopulation penetrance between MGE types, with IS_Tn and Phage particularly prone to varying across individuals within a host population (**Extended Data fig. 11D**). Because MGE sizes vary and read mapping for lesser-sampled taxa is noisy, these numbers represent conservative lower bounds on active MGE penetrance into cellular population genomes. Even so, the extent of partial carriage observed highlights MGEs’ role in what amounts to large ongoing *in situ* knockout experiments: among 372 IS_Tn elements that both interrupt a protein-coding gene and were amenable to partial carriage analysis, 81 (∼22%) variably interrupt or leave intact host protein-coding genes. Together, these results suggest that MGE penetrance within member genomes of any given population differentially provide individuals access to genetic diversity, while shielding the population from loss of function due to MGE insertion (*59, 60*).

## Conclusions

MGEs have long been known from cultivated isolate genomes to be important drivers of strain-level variation and key functional adaptations, particularly in biomedical contexts related to AMR (*22, 23, 32*). However, virtually no data exist for diverse MGE types in natural settings. Our analyses presented here for Stordalen Mire microbiomes illuminate the mechanisms and scope of MGE impact on a climate-sensitive ecosystem by showing that, well beyond biotic interactions, the mobilome affects functions intrinsic to a wide range of physiologically and ecologically significant processes. Strikingly, the IS_Tn elements that heavily dominate recovered MGE recombinases in our dataset are also the most active by two of our three measures (i.e., transcription and genomic neighborhood change) (**Fig. 5B**), highlighting the need for future studies to examine context-dependent MGE effects like modulation of host gene expression downstream of MGE insertion sites.

As improved methods for studying the genomic potential of microbes in nature come on-line, the field has progressed from community-level taxonomic and functional descriptions towards species-resolved links between taxa and their functions (*28, 34*). Continued advances in sequencing and analytical approaches are increasing this resolution to reveal intra-population variation, early speciation, and selection (*8*) with recent MGE typing efforts (*32*) and further analytical advances made here finally bringing a quantitative understanding of the mobilome within reach, even in complex natural systems. This work benchmarks sequencing artifacts and provides empirical baselines and inferred impacts for all major MGEs in a natural soil system. These findings suggest that ∼1% of MGE recombinases are active, genomes contain 36–66 MGE recombinases on average, and that MGEs impact (via carriage or interruption) diverse ecosystem-relevant functions. As technologies advance, the mobilome will come more fully into focus, and, given this new toolkit and baseline findings, we predict it will help reveal a subtler layer of change that could serve as the canary in the coalmine for early detection of change.

## Acknowledgments

Non-author contribution: we thank Ohio Supercomputer Center for providing computing resources.

## Author contribution

J.G., S.R., M.B.S., and S.C.B. conceptualized, designed, coordinated the study. S.A., J.G. performed long read and short read comparison. S.A., A.F., J.G. performed MGE recombinase host lineage analyses. G.D.-H., D.S., D.V., S.C.B., A.A.P., G.S., F.T., C.H.-V., S.S., Z.Z., C.O.A., J.G. performed MGE impacted function analyses. D.C., S.A., S.B.H. performed metagenome assembly and binning. J.G., S.R., M.B.S., and S.C.B. drafted the manuscript with contribution from B.J.W., G.W.T., V.I.R. All authors read, edited, commented on, and approved the final manuscript.

## Data and material availability

The authors declare that all the data in this study are freely and publicly available without restriction. The metagenomic and metatranscriptomic raw reads are available in SRA / ENA under Umbrella BioProject PRJNA888099 with individual accessions cited in **table S1**. The data provided by the DOE Joint Genome Institute were generated under proposals “10.46936/10.25585/60001148” and “10.46936/fics.proj.2017.49950/60006215”. MAGs used in this study are available in Zenodo with https://doi.org/10.5281/zenodo.13984630 (main set of 36,419 MAGs) and https://doi.org/10.5281/zenodo.14207415 (733 short-read, long-read, and hybrid MAGs from six paired samples). The paired long read and short read data from six samples are available in PRJNA888099 with accessions cited in **table S7**. Processed data including tables for MGE recombinase identification, activity by metatranscriptome, mobility by genomic neighborhood change, and partial carriage, and MGE impacted functions are available through supplementary tables and a large file data package shared through CERN’s Zenodo repository (https://zenodo.org/records/15102623). The code produced in this study can be found in the project’s GitHub repository (https://github.com/emerge-bii/mobilome-submit-202407).

## Supplementary Materials

### Methods and Materials

#### Data collection, metagenome assembly and binning, gene annotation and taxonomy assignment

Our study site, Stordalen Mire, is a long-term ecological research site in northern Sweden. Large sampling and sequencing efforts since 2010 have yielded ∼4.9 Tb of metagenomic sequence data from 615 readsets from 408 bulk soil samples that span space (across the site’s three habitat types, n = 188 palsa, 200 bog, 227 fen; average ∼8 Gb/sample) and time (summer peak growing season, 2010–2017) (**table S1**) with details described in another parallel study (*31*) and previously (*28, 29, 61*). The IsoGenie Consortium and EMERGE Biology Integration Institute have developed extensive resources from Stordalen Mire metagenome data, including assembly across all samples to produce 30,710,180 contigs (≥1500 bp) and 36,419 medium to high-quality metagenome-assembled genomes (MAGs; completeness ≥ 70% and contamination ≤10%, CheckM2 v1.0.2) (*62*) that clustered into 2,048 populations (95% Average Nucleotide Identity [ANI]) per community standards (*63*). Select samples have also been sequenced using metatranscriptomics to assess gene expression (50 spatio-temporally diverse samples; average ∼18 Gb per sample). For this work, we additionally performed paired long- and short-read DNA sequencing on six samples, chosen to be representative of the larger dataset, as a control to assess short-read MGE recovery.

The protocol used for sample preparation and sequencing for short-read metagenomes has been described previously (*28, 29, 61*). Briefly, nucleic acids were extracted according to a published protocol (*64*), except for the MAG source samples from substrate addition incubations (“201607_SubstrateInc_9to19”), which were extracted with the DNeasy PowerSoil Kit (cat# 12988-100, Qiagen). Libraries were prepared with either KAPA Hyperprep, TruSeq, or Nextera XT kits for short-read metagenomes; SMRTbell Express Template Prep Kit 2.0 for long-read metagenomes; or TruSeq Stranded mRNA kit for metatranscriptomes. Reads were sequenced with either Illumina HiSeq (2000 or 2500), NovaSeq 6000, or NextSeq 500 for short-read metagenomes; PacBio Sequel IIe for long-read metagenomes; or NovaSeq 6000 for metatranscriptomes.

Short-read sequencing data from individual samples were trimmed by Trimmomatic (v.0.36) (*65*), assembled by SPAdes with --meta option (v.3.12) (*66*), and then binned by MetaBAT2 (*67*), GroopM2 (*68*), and MaxBin2 (*69*) (Cronin.v2), or an ensemble approach (*31*). Those sequenced at JGI were processed by JGI’s standard assembly and binning pipeline. Briefly, reads were quality-controlled, filtered, and trimmed using the BBTools package (*70*), assembled with SPAdes with --meta option (*66*), and binned with MetaBAT2 (*67*). The bins produced from this effort were combined with 9,406 bins from Woodcroft 2018 (*28*) and 11,120 bins from stable isotope probing (SIP) experiments with field peat soil and labeled litter (sequenced with NovaSeq at JGI and assembled with MEGAHIT v1.1.3) (*71*), and then filtered using 70% completeness and 10% contamination CheckM2 (v1.0.2) (*62*) cutoffs to form the 36,419 metagenome-assembled genomes (MAGs; consisting of 4,193 from Cronin.v1, 27,423 from Cronin.v2, 847 from JGI, 1,543 from Woodcroft 2018 and 2,413 from SIP). Next, bins obtained from the same metagenome were de-duplicated using Galah (95% ANI, commit e9e59a8) (*63*), and all bins across the entire dataset were further dereplicated at 95% ANI using Galah (commit e9e59a8), producing 2,048 population level clusters. Further, the contigs encoding a recombinase were annotated with DRAM (--min_contig_size 1500) (*72*) and taxonomically assigned at phylum level with CAT (*73*) requiring a minimal contig length of 3kb. The contigs in MAGs were assigned to the MAG taxon (obtained with GTDB-tk) (*74*), since taxonomic assignments based on whole MAGs are typically more reliable than single-contig assignments.

#### MGE recombinase identification

Recombinase identification was based on a framework recently developed to robustly identify and classify MGE recombinases from microbial isolate genomes (*32*). Here, we applied the same recombinase HMMs profiles and suggested cut-offs to identify MGEs in metagenomic assemblies. In brief, open reading frames were predicted from assembled contigs by prodigal (v2.6.3) using the “-p meta” option (*75*), then recombinases were identified among the predicted protein sequences by hmmsearch (v3.3.2) (*76*) using the 68 recombinase subfamily HMMs (**table S2**) and the “--cut-ga” option. Identified recombinases were assigned to the respective MGE recombinase type corresponding to the HMM with best score – as suggested previously (*32*).

#### Annotation-based inclusion/exclusion criteria

MGE recombinase candidates were further quality controlled as recommended previously (*32*) by using eggnog-mapper (v2.1.9, default settings) (*77*). First, genes annotated as invertase, replication protein, DNA repair, DNA binding, homologous recombination, viral exonucleases, or reverse transcriptase (interpreted to be spurious hits to other host genes related to DNA manipulation) were removed, except for the ones also annotated with the Pfam catalytic domains of “transposase” or “helicase” that were retained after manual curation. This removed 0.68% of the identified possible recombinases (**Extended Data fig. 1**). Second, sequences annotated by eggnog-mapper as CRISPR, defense response to virus, or maintenance of DNA repeat elements were removed to filter out spurious hits thought to be similar to Cas family recombinases, but not considered as an MGE recombinase. This removed 0.46% of the identified possible recombinases. Third, we discarded predicted recombinases that contained only an HTH (DNA binding) domain, as well as relaxase hits for which Pfam annotates a helix domain, but without the catalytic domain required for recombinase activity. This removed 0.07% of the identified possible recombinases. Fourth, recombinases matching a “Cell” type recombinase HMMs (and thus unlikely to be encoded by an MGE) were removed. This removed 2.52% of the identified candidate recombinases, and yielded a final total of 2,107,204 MGE recombinases (**table S3** item A). Lastly, some recombinase subfamilies/HMMs include recombinases from different MGE types, which prevents assignment for that HMM to a specific MGE type. These recombinases with ambiguous origin accounted for 20.51% of the identified recombinases. They were *included* in the analyses of total MGE recombinases, but *excluded* in analyses of individual MGE types, which focused on four recombinase-based MGE types: “IS_Tn” (insertion sequence [IS] and transposon [Tn]), “CE” (conjugative elements including conjugative element and mobility islands), “PhL” (phage and phage-like elements) and “Integron” (integron) (**Extended Data fig. 1**). To assess the diversity of these these MGE recombinases, we considered recombinase sequences as “redundant” when they clustered with other sequences at 100% AAI using CD-HIT (v4.8.1) (*78*) with “-c 1 -g 0 -G 1 -l 30 -d 0 -M 0”. This yielded 1,130,776 clusters that we considered as unique recombinases.

### Identification of cargo genes/interrupted genes associated with MGEs

Once MGE recombinases were identified, we surveyed their genomic context to assess potential cargo and/or interrupted genes. This was done using MGE-specific approaches as follows.

*Integron*: All integron-containing contigs were screened for cargo genes (also known as “gene cassettes”) using integron-finder (v2.0.2) (*79*) with default parameters. Briefly, integron-finder identifies integrons by using integron specific integrase HMMs to identify integron integrases (*intI*), and a covariance model to identify the integron attachment site (*attC)*, as the two bounding regions of an integron sequence. Only integrons encoding the *intI* and at least one *attC* treated as “complete” by integron-finder were retained. There are two types of integron structures that are known in terms of the orientation of *intI* and *attC*: 1) a regular type with *intI* and *attC* on opposite strands and facing outwards, 2) another type with *intI* and *attC* on the same strand, called inverted integrase integron (*79*). Of the 686 “complete” integrons we identified, 681 integrons also met these orientation expectations and the 5 that did not were not considered further. The gene cassettes delimited by *intI* and *attCs* in these integrons were treated as cargo genes associated with integrons (*80*), resulting in integron regions with various lengths (minimum: 983bp, median: 2,310bp, mean: 3,092bp, and maximum: 14,195bp). Problematically, most contigs (97.4% or 25,953 out of 26,634) encoding Khedkar’s “Integron” type recombinase were not considered complete by integron-finder. To test this, we also replaced the integron integrase HMM in the integron-finder tool with that from Khedkar et al. (*32*) and found it produced nearly identical results and did not yield additional instances of predicted complete integrons. One explanation could be that *attC*, not integron integrase, is limitation, since the flanking *attC* sequences are identified using a database of known *attC* sequences and these could miss novel *attC* sequences used by integrons in natural microbiomes. Given this, we interpret the large discrepancy between Khedkar’s and integron-finder’s integrons to represent one of the following scenarios: (i) degraded or domesticated integrons that lack *attC* sites (*81, 82*), (ii) integrons that lacks *attC* sites due to fragmented assembly, or (iii) intact and likely complete integrons that have novel *attC* sites. Thus our analyses will under-estimate the extent of cargo genes carried by integrons in Stordalen Mire.

*Conjugative Element (CE):* CE are made of three key components – a relaxosome (includes the CE-type recombinase), a conjugation couple protein, and a type IV secretion system (T4SS) – that can be used as hallmark genes that are more conserved than other CE genes (*83*). To identify cargo genes associated with CE, the CE hallmark genes were identified in CE recombinase-encoding contigs by conjscan implemented in MacSyFinder (v2.0rc7) with “--models CONJScan/Plasmids all --db-type gembase” and “--models CONJScan/Chromosome all --db-type gembase” with gene protein sequences as input (*83, 84*). Conjscan specifically establishes 8 types of CEs using hallmark genes that are core across all 8 CE types, and accessory genes that are diagnostic for specific CE types. CE identified by conjscan from Stordalen Mire sequence data were those that matched a relaxase, a couple protein (an ATPase that couples MGE with the T4SS), a T4SS ATPase, and an additional structural gene from T4SS. For the purposes of identifying cargo genes, matches of one model, MOB, were removed since it only requires the relaxase and thus does not identify the boundaries to establish a complete CE. As a result, we identified 1,017 near complete CEs. As recommended, we required the distance between the hallmark and accessory genes to be <=30 genes except for the relaxase which could be as much as 60 genes away from any other hallmark and accessory genes (*83*). In a small number (13 of 1,017) of instances conjscan identified >1 CE in a region. For these, we manually separated the identified hallmark and accessory genes into subgroups according to gene proximity and required each subgroup to include all hallmark genes (relaxase, conjugation couple protein and T4SS ATPase) and at least one accessory T4SS gene. Once such “complete” CEs were identified, the genes between the leftmost and rightmost hallmark and accessory genes were assumed to be cargo genes carried by the CE (except for hallmark and accessory genes themselves). This resulted in CE regions with various lengths (minimum: 7907, median: 16,290, mean: 23,383, and maximum: 81,148 bp). Overall, 1.6% of the 62,660 CE type recombinase genes identified by our analyses were considered near-complete CE by conjscan. Again, this was not a function of improved HMMs as we tested replacing the relaxase HMMs in conjscan by those from Khedkar et al. (*32*) and found it produced similar results. Instead we interpret this large discrepancy to suggest that most CEs in these data are degraded, fragmented from poor assembly, or can not be formally detected as complete because of limitation and bias in hallmark gene databases.

*Insertion sequence / Transposon:* To identify “IS_Tn” recombinase interrupted genes, up to 3,000 bp upstream and downstream of “IS_Tn” recombinases were collected from each contig, requiring a minimal length of at least 50 bp on each end and at least 500 bp across the two ends combined (1,063,800 IS_Tn recombinases in total meeting the requirements), and then aligned by minimap2 (v.2.22, default settings) (*85*) to the coding sequence of representatives of protein clusters (PCs) obtained by MMseqs2 at 95% identity (v4.8.1) (*86*) from all contigs (≥1500 bp) used in this study. The corresponding coding sequence was considered interrupted if the following four conditions were met: 1) upstream and downstream alignments combined covered ≥50% of the coding sequence of the reference PC; 2) if there is a gap between upstream and downstream alignments of the coding sequence, the gap is ≤10% of the coding sequence length; 3) if there is an overlap between upstream and downstream alignments on the coding sequence, the overlap is ≤10% of the shorter alignment; and 4) if there are multiple alignments in either upstream or downstream to the coding sequence, the pair with largest coverage of coding sequence is chosen. This identified 5,342 PCs putatively interrupted by a total of 12,332 IS_Tn type recombinases.

Phage / phage-like elements: All contigs encoding “Phage” recombinase were screened for temperate phage by VirSorter2 (v2.2.3) following VS 2 SOP step 1 (v3.0) (87, 88), and the predicted phage sequences trimmed by checkV (default setting, version 0.7.0) (89). Next, the checkV trimmed sequences encoding a “Phage” recombinase were considered as phage/phage-like regions. To identify cargo genes, i.e. host(-like) genes encoded on phage/phage-like elements, we opted for a conservative approach in which we reduced the dataset for “cargo gene inspection” to include only phage/phage-like regions with one “Phage” recombinase, and use the leftmost and rightmost checkV called “viral gene” or “Phage” recombinases as the boundaries. This identified 4,277 temperate phages with “Phage” recombinases and at least one extra gene other than two at ends. Meanwhile, 347 phage/phage-like elements were removed for having more than one “Phage” recombinase, in order to avoid cases where multiple contiguous temperate phages or degraded ones yielded a single prophage prediction, which could lead to host genes being mistaken as cargo genes. This resulted in 3930 conserved phage/phage-like regions (minimum: 1386bp, median: 7293bp, mean: 13,779bp, and maximum: 352,302bp).

*Annotations of cargo/interrupted genes:* Insertion sequence or transposon interrupted genes and genes within other MGEs were functionally annotated using KEGG and Pfam databases in DRAMv for phages/phage-like elements or DRAM for others MGEs (v1.2.3, -- min_contig_size 1500) (*72*), and further evaluated by comparison with best hits in NCBI NR (downloaded on 10/04/2020) using DIAMOND (v2.0.8.146, blastp bit-score ≥50) (*90*). A subset of the genes of specific interest to the Stordalen Mire thawing permafrost system were selected and manually curated for display (**table S4**) including Transport, Carbon/Nitrogen/Sulfur cycling, Signaling, Antiviral defense, and Genome maintenance and expression.

#### Evaluation of short read assembly and binning artifacts for recovering MGE recombinases using long reads

*Assembly and binning of paired long reads and short reads*: Soil samples (6 total) were collected at the Stordalen Mire site from 2019 from two depths (cm below ground) across three habitats (Palsa, Bog, and Fen). DNA extraction and Illumina NovaSeq sequencing was performed as described above. In addition, DNA sequencing was performed from the same DNA extractions using PacBio Sequel IIe HiFi sequencing platform at JGI. PacBio data was processed at JGI to form filtered CCS (Circular Consensus Sequencing) reads. Assemblies were generated through two methods: short-only and hybrid short-read plus long-read (herein: hybrid). Short-only was assembled with metaSPAdes v3.15.4 (*66*) using Aviary v0.5.3 (*91*) with default parameters. Hybrid assembly was performed using Aviary v0.5.3 with default parameters. This involved a step-down procedure with long-read assembly through metaFlye v2.9-b1768 (*92*), followed by short-read polishing by Racon v1.4.3 (*93*), Pilon v1.24 (*94*) and then Racon again. Next, reads that didn’t map to high-quality metaFlye contigs were hybrid assembled with metaSPAdes v3.15.4 and binned out with MetaBAT2 v2.1.5 (*67*). For each bin, the reads within the bin were hybrid assembled using Unicycler v0.4.8 (*95*). The high-coverage metaFlye contigs and Unicycler contigs were then combined to form the assembly fasta file. Genome recovery was performed using Aviary v0.5.3 with samples chosen for differential abundance binning by Bin Chicken v0.4.2 (*96*) using SingleM metapackage S3.0.5 (*97*). This involved initial read mapping through CoverM v0.6.1 (*98*) using minimap2 v2.18 (*85*) and binning by MetaBAT v2.1.5 (*7*), MetaBAT2 v2.1.5 (*67*), VAMB v3.0.2 (*99*), SemiBin v1.3.1 (*100*), Rosella v0.4.2 (*101*), CONCOCT v1.1.0 (*102*) and MaxBin2 v2.2.7 (*69*). Genomes were analyzed using CheckM2 v1.0.2 (*62*) and clustered at 95% ANI using Galah v0.4.0 (*63*).

*Assessment of MGE recombinase recovery by short-read assembly and binning*: We considered the high quality hybrid MAGs as the best available references and used these to assess the recovery of MGE recombinases in short-read contigs and MAGs. A total of 17 high-quality MAGs (completeness ≥95%, contaminant ≤5% and at most 100 contigs) from hybrid assembly were selected and paired with 17 MAGs recovered from short-read assembly through the 95% ANI clusters (the highest ANI were chosen if there were many) for comparison. The short-read MAGs included 5 medium quality (≥70% completeness, ≤10% contamination) and 12 high quality (≥95% completeness, ≤5% contamination).

Evaluation of these 17 MAGs were carried out as follows. First, each pair of hybrid MAG and short-read MAG were compared in terms of total number of MGE recombinases, as well as unique and shared MGE recombinases at 100% AAI by CD-HIT (v4.8.1) with “-c 1 -g 0 -G 1 -l 30 -d 0 -M 0” (78). Second, we evaluated the impact of sequencing depth, assembly, and binning to assess which step was most problematically leading to under-detection of MGE recombinase identification in short reads. Initially, we mapped short-read assembled contigs (sheared into 500 bp fragments) to hybrid MAGs from the same sample using coverM v0.6.1 requiring each read to be 75% aligned with 95% identity (98). Third, these data were used to assign MGE recombinases in the 17 hybrid MAGs into one of three categories: (i) those recovered by short-read MAGs (termed “covered”), (ii) those not fully covered by short-read contigs, ie, missed by short-read assembly (termed “missed assembly”), and (iii) those fully covered by short-read contigs, ie, recovered by short read assembly but missed by short-read binning (termed “missed binning”). Fourth, the coverage of each gene was generated by dirseq v0.4.3 (28) using the BAM file from above and used to assess the relative abundances of each category as a metric for the impact on MGE-representation in the short-read assemblies.

With a better understanding for which step was failing most often for MGE recombinase recovery in short-read assemblies, we sought next to evaluate what factor(s) might be causing the failed capture events. First, we evaluated sequencing depth since low coverage might cause breakages in short-read assembly graphs. This was done by mapping trimmed short reads to hybrid MAGs using coverM v0.6.1 requiring each read 75% aligned with 95% identity, calculating the read coverage of each gene by dirseq v0.4.3, and comparing coverages among the three categories of MGE recombinases. Second, we compared the genomic neighborhood number among the three categories, reasoning that perhaps indels were an issue. Genomic neighborhood number was calculated by clustering genomic neighborhoods of each unique MGE recombinase (see below) at 90% ANI. To test this, nucleotide diversity for each base position was calculated as 1 − [(number of “A” bases / total bases) ^2 + (number of “C” bases / total bases) ^2 + (number of “T” bases / total bases) ^2 + (number of “G” bases / total bases) ^2 + (number of “Deteletion” bases / total bases) ^2 + (number of “Insertion” bases / total bases) ^2], adopting the nucleotide diversity definition from inStrain (*8*) with addition of deletion and insertion variants. The variant count at the base position was generated by pysamstats (v1.1.2, https://github.com/alimanfoo/pysamstats) with “--type=variation” option. Third, we assessed nucleotide diversity of each MGE recombinase, reasoning that perhaps elevated nucleotide diversity could lead to short-read assembly graph breakages. To test this, for each of the 17 MAG pairs, MGE recombinases only identified in hybrid MAG were compared to short-read assembled contigs to track whether they were assembled by short reads and then whether they were binned in MAGs from other MAG clusters. The tracking of MGE recombinases in contigs was accomplished with “MMseqs map” (v13.45111) with default setting and then filtered for hits with 100% identity in protein sequence (*86*).

#### MGE recombinase per genome estimation from contigs and MAGs

To estimate MGE recombinase number per genome from contigs, the median count of ribosomal genes was first used to estimate the number of genomes in each assembly. To do this, ribosomal genes were extracted from contig annotations by DRAM (*72*) using KEGG IDs (table S6), and the average number of recombinase genes per genome was calculated by dividing the total number of recombinases in the sample by the estimated total number of genomes. We next linked individual recombinase-encoding contigs to MAGs built from the same sample. To that end, we used exact string searching to identify MGE-recombinase-encoding contigs in the de-replicated MAGs from the same sample. Through this process, 106,017 out of a total 665,700 (15.9%) MGE-recombinases-encoding contigs ≥3kbp could be linked to a MAG. To account for partial MAG recovery, the number of MGE recombinases per genome estimated from MAGs was calculated as the number of MGE recombinases in a given MAG divided by MAG completeness.

#### Genomic neighborhoods of MGE recombinases and mobility evaluation

MGE mobility was assessed by comparing the genomic neighborhoods of individual recombinases. For this analysis, we limited the input data to recombinases that occurred ≥2 times in the total dataset (assembled contigs per sample). This was achieved by clustering the recombinases at 100% AAI using CD-HIT v4.8.1 with “-c 1 -g 0 -G 1 -l 30 -d 0 -M 0” (*78*), only keeping the clusters with at least two recombinases. For each MGE recombinase, 300 bp upstream and downstream of the recombinase were considered as its immediate genomic neighborhoods, excluding those near the contig ends (< 300bp). For each recombinase cluster, pairwise comparison of neighborhoods was performed among all member recombinases using both neighborhoods (ie, upstream vs. upstream, and downstream vs. downstream), using BLAST+ (v2.8.1+) with “-task blastn” (*103*) and bitscore ≥50. Average nucleotide identity (ANI) for each neighborhood comparison was calculated as the “sequence identity from blastn output” multiplied by the “alignment coverages’’. For each comparison, we labeled the neighborhood with the higher ANI value as “high_ani” and the neighborhood with lower ANI as “low_ani”. For some MGEs, we expect the neighborhood on the MGE side of the recombinase to be part of the MGE and thus remain identical after integration into a new locus, so we used “low_ani” values (i.e. the lower ANI between the upstream and downstream region of the pair) to evaluate whether the neighborhood of an MGE changed or not. An MGE recombinase pair was considered to be in different genomic locations if “low_ani” was <1%, and a recombinase cluster was treated as active if one pairwise comparisons showed different neighborhoods. This approach only used immediate neighborhood (within 300bp) since a requirement of neighborhood with longer range would remove more recombinases from this analysis due to the short average contig length. However, as a sensitivity analysis, we also tested the above with 600 bp and 1000 bp neighborhoods. These analyses showed that the size of the genomic neighborhood (300 - 1000bp) had very little impact on the fraction of MGE recombinases identified as mobile via this changing genomic neighborhood metric (**Extended Data fig. 10**).

#### MGE recombinase activity by metatranscriptome

MGE activity was assessed via documenting which MGE recombinases were detectably transcribed in metatranscriptomes. Leveraging 81 paired metatranscriptomes and metagenomes, we evaluated MGE recombinase transcriptional activity by mapping metatranscriptome reads to contigs assembled from metagenomes using transcriptM v0.3.1 (*104*) with ORF predictions from DRAM (see above) as the “--gff” input, which also removed tRNA, rRNA, tmRNA, and PhiX from the reads. To ensure the quality of RNAseq data (reverse stranded), we only analyzed 58 pairs with ≥80% of reads mapped to the coding reverse strand of ORFs in contigs, based on dirseq v0.4.3 (*28*) with BAM and GFF files from above as inputs (table S5). We only considered contigs ≥10 kbp to avoid spurious ORF predictions from short contigs. Next we defined genes with ≥90% of gene length covered as expressed using pydirseq v0.1.0-alpha with BAM and GFF files from above as inputs (*105*). To normalize for the differences in sequencing depth and overall activity of the microbial community across samples, we used the average percentage of ribosomal genes (table S6) detected as active as the baseline activity to normalize other genes (termed “normalized active ratio”), since ribosomal genes are expected to be constitutively expressed in active cells. This “normalized active ratio” thus provides an estimation of the number of genes detected as expressed within a functional category (e.g. recombinases) relative to an estimated total number of genomes detected as transcriptionally active in the same sample.

#### MGE recombinase partial carriage by host population

As a result of MGE movement within the host population and an equilibrium of the co-evolution of MGEs and their hosts (*106*), we noticed that some MGE recombinases were present in only a fraction of the contigs assigned to a given host population (termed “partial carriage”). To quantitatively evaluate this, trimmed metagenomic reads were mapped to contigs from the same sample using coverM v0.6.1 with default setting (*98*), then the resulting BAM file was filtered requiring ≥90% each read aligned with ≥95% identity, and the filtered BAM file was used as input to “samtools depth” (v1.17) with “-a” option to get the read depth coverage at each position (*107*). The first and last 150bp of each contig were not considered since read depth coverage near the ends are expected to be artificially lower. We defined partial carriage as cases where the coverage of the MGE recombinase was at least one standard deviation lower than the larger one (requiring a minimal coverage of 1x) of the average coverages of the upstream and downstream contig regions (Cohen’s d ≥1). To ensure enough host regions were considered for reliable mapping-based estimation of genome coverage, the MGE recombinase total set was filtered to 23,647 MGE recombinases with ≥20 kb upstream and downstream. Although, again sensitivity analyses were conducted, and similar patterns were observed when analyzing 2.5 kbp instead of 20 kbp regions (Extended Data fig. 11), which suggests that this conservative cutoff did not substantially influence the detection of partial carriage in these data.

We expect that this method should give us a lower-bound estimate of partial carriage, since it is not able to detect cases for which (i) the hosts have low read coverage (rare), (ii) the carriage ratio is lower than but close to 1, (iii) host genome coverage has a wide range of variation in different genomic locations due to strain variation within the population, or (iv) the MGE recombinase is on the edge of a contig.

#### Statistics

All statistics and visualization were performed in R v4.2.1 (*108*). The R packages and versions used can be found in https://github.com/emerge-bii/mobilome-submit-202407. For statistical significance between groups, Wilcoxon rank sum test was used for two groups and Kruskal Wallis test was used for >2 groups as implemented in “compare_means” in ggpubr package v0.6.0, with *P values* adjusted by Benjamini-Hochberg adjustment for multiple comparisons.

## Extended Data

**Extended Data figure 1.**
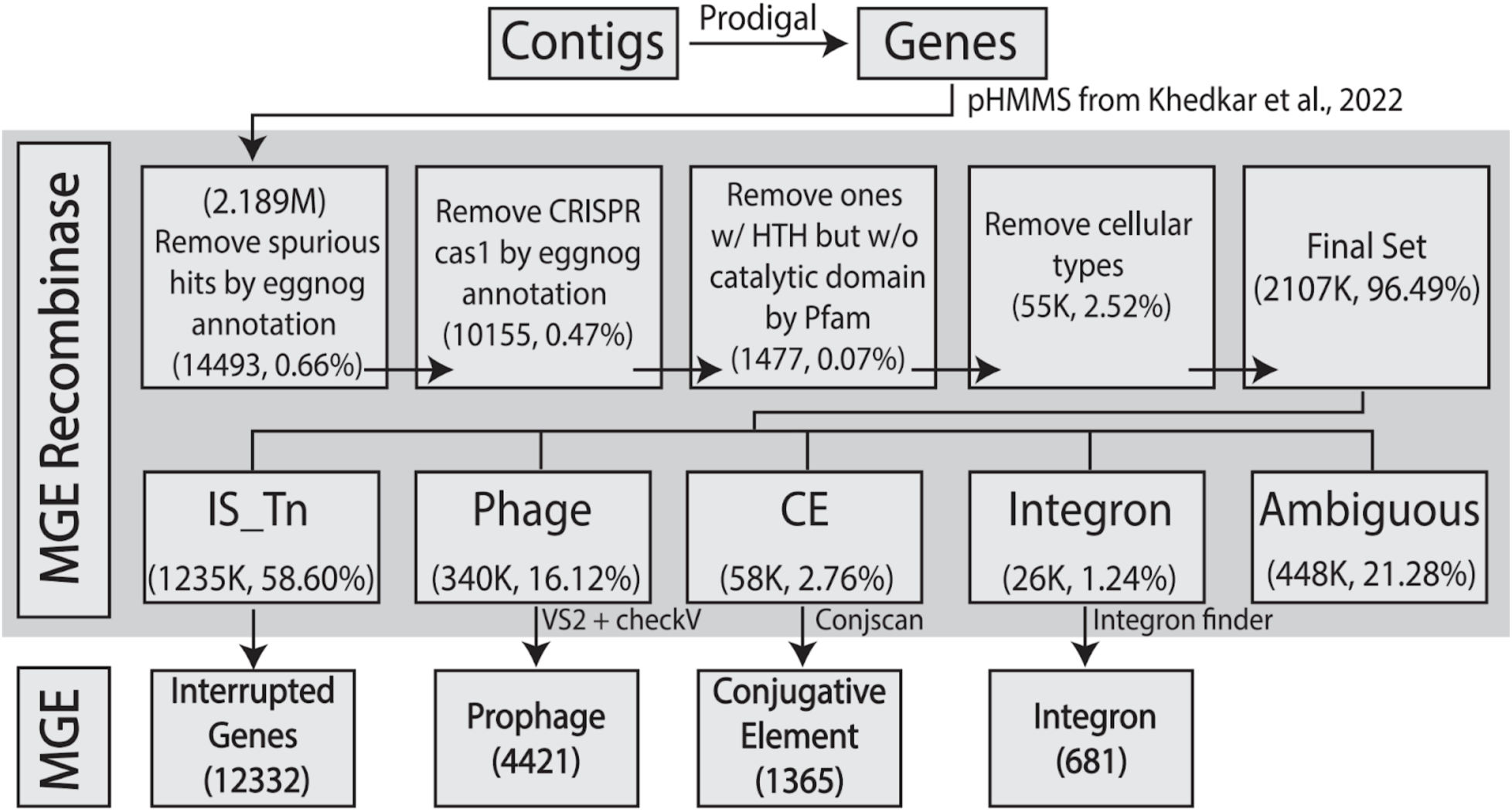
Flowchart of MGE recombinase and MGE boundary identification. A total of ∼2.2M recombinases were identified using 68 HMMs, filtered in four steps, and then the resultant ∼2.1M MGE recombinases were classified into ‘types’ according to their HMM affiliation using criteria established previously (*18*). These identification and typing steps are collectively denoted by the background gray shading. The numbers and percentages of recombinases involved in each step are shown in each box. Within each MGE recombinase type, below the gray shaded area, various strategies to conservatively identify boundaries were used (see tools listed in figure) largely by using hallmark genes annotated by these tools as conservative boundaries (see Methods). In the remainder of the study, the ∼2.1M MGE recombinases identified were used for host lineage and activity/mobility inferences, whereas only the boundary-constrained used for MGE-impacted metabolism analyses.

**Extended Data figure 2.**
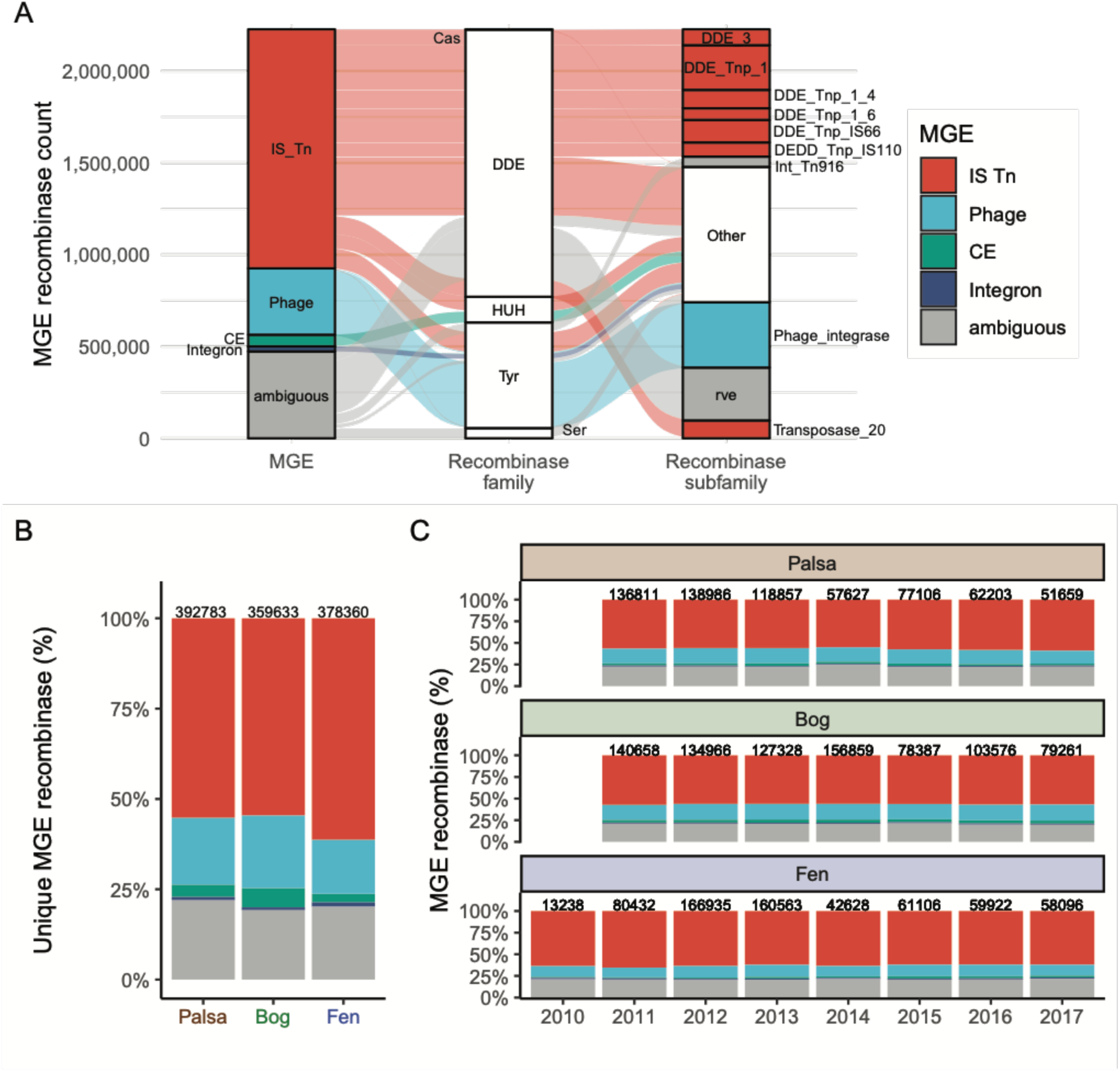
Overview of MGE recombinase distribution from Stordalen Mire. **A.** Distribution of contig-identified MGE recombinases by MGE type, as used in the main text, as well as MGE recombinase family and subfamily (see a full list described in **table S2**). **B.** Unique MGE recombinase (clustered by 100% AAI) by habitat. The numbers at the top of bars are the number of unique MGE recombinases. **C.** Total MGE recombinase distribution by habitat and year. The numbers at the top of bars are the number of total MGE recombinases.

**Extended Data figure 3.**
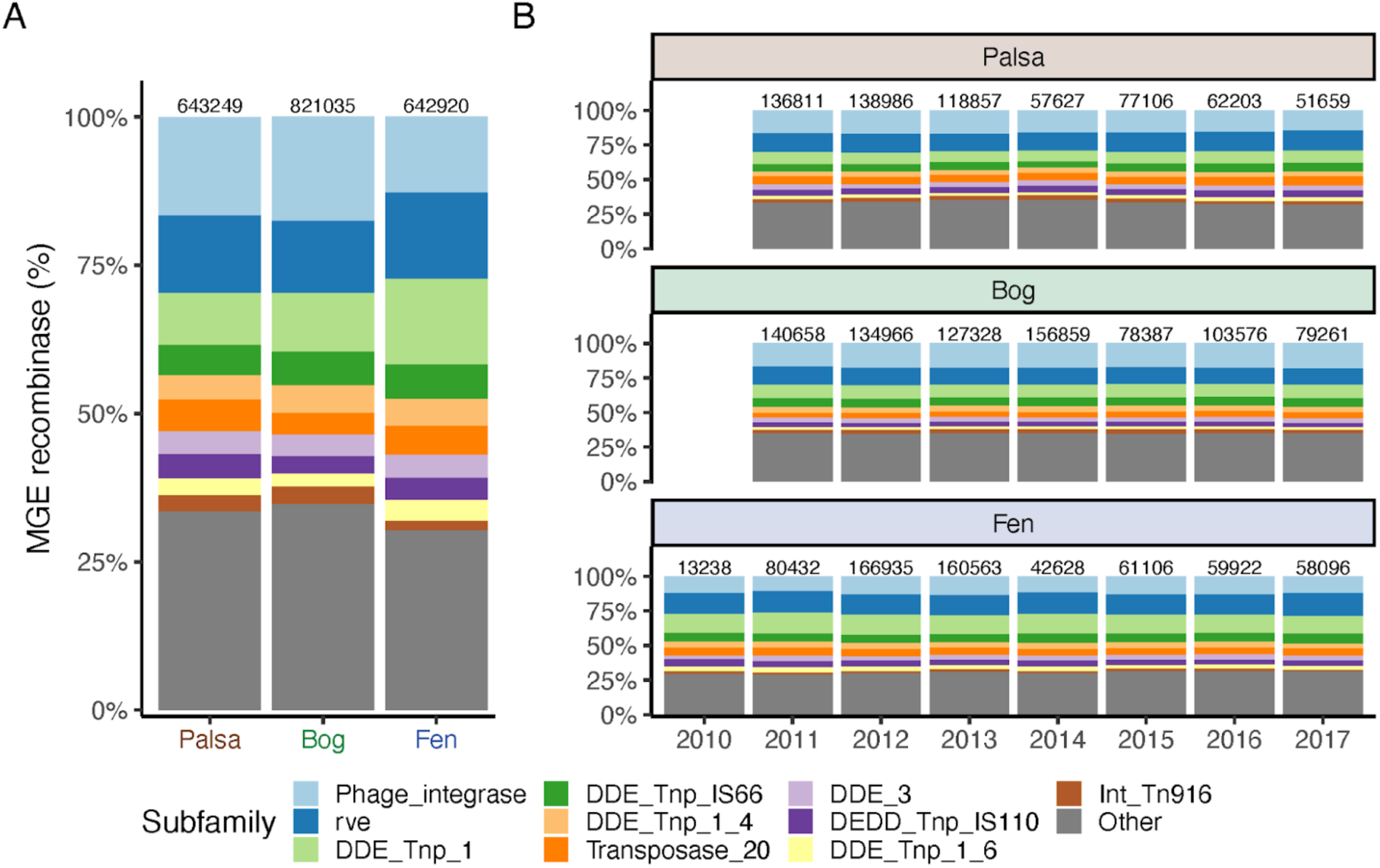
MGE recombinase distribution based on subfamily. **A.** MGE recombinase subfamily distribution by habitat. The numbers at the top of bars are the number of total MGE recombinases. **B.** MGE recombinase subfamily distribution by habitat and year.

**Extended Data figure 4.**
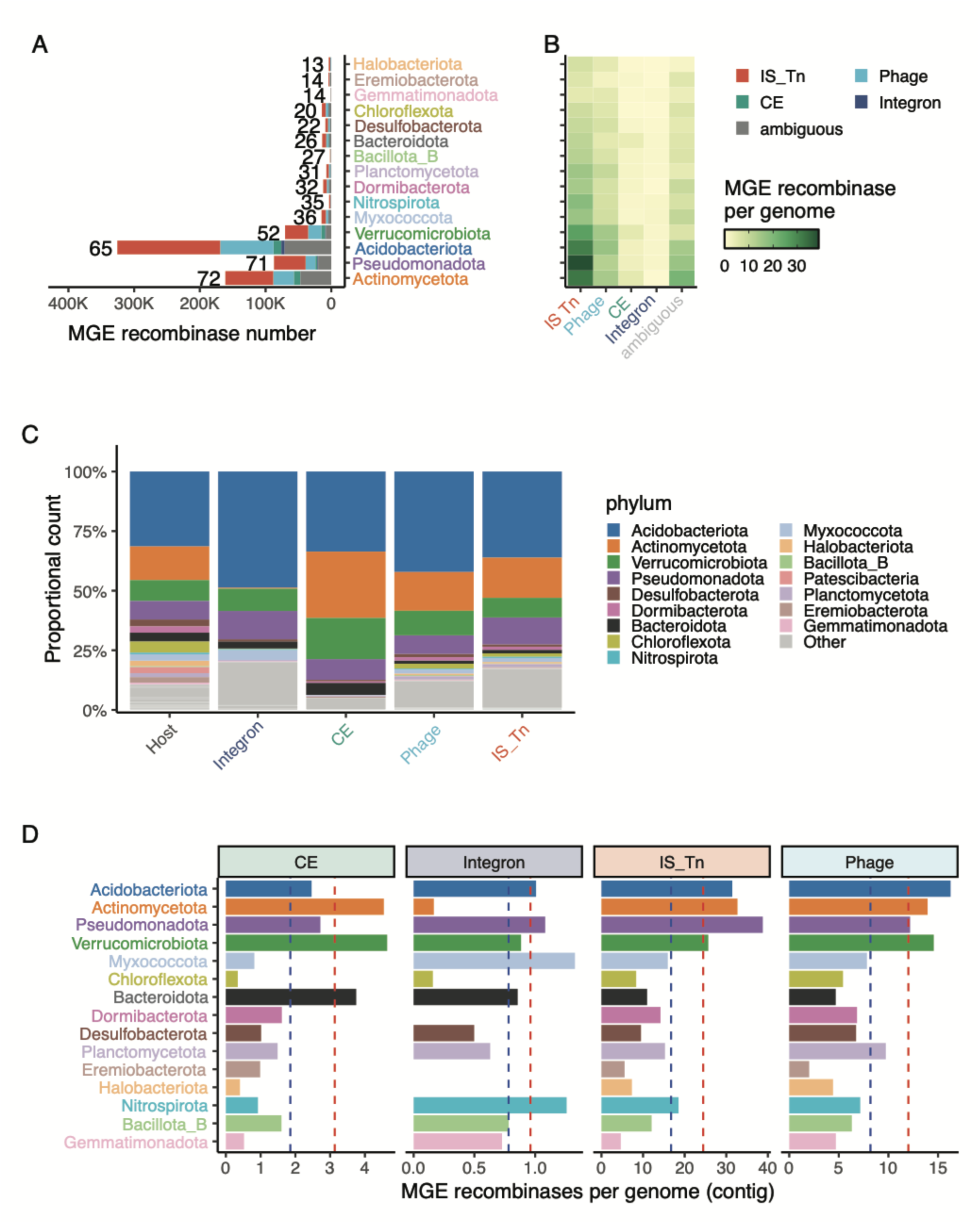
MGE host lineage based on MGE recombinase identified from contigs. MGE recombinase encoding contigs (>=3kb) were evaluated for taxonomic patterns as a complement to MAG-based analyses in Figure 3. **A.** Number of each MGE type found in each phyla where the top 15 phyla (out of 89 in total) accounting for 84.4% of total MGE recombinases are shown, ordered by MGE recombinase count per genome (the number to the left of each bar).The genome number is estimated from ribosomal genes (see Methods). Phyla names are colored per Figure 3. **B.** The number of MGE recombinases identified per genome grouped by MGE type and host phyla. **C**. Phylum distribution of the overall microbial community (estimated from ribosomal gene encoding contigs) and MGE recombinase types. **D**. MGE recombinase number per genome per phylum. The dashed red lines are the average MGE recombinase number per genome across all phyla, and the blue dashed lines are the average across the phyla shown in the plot.

**Extended Data figure 5.**
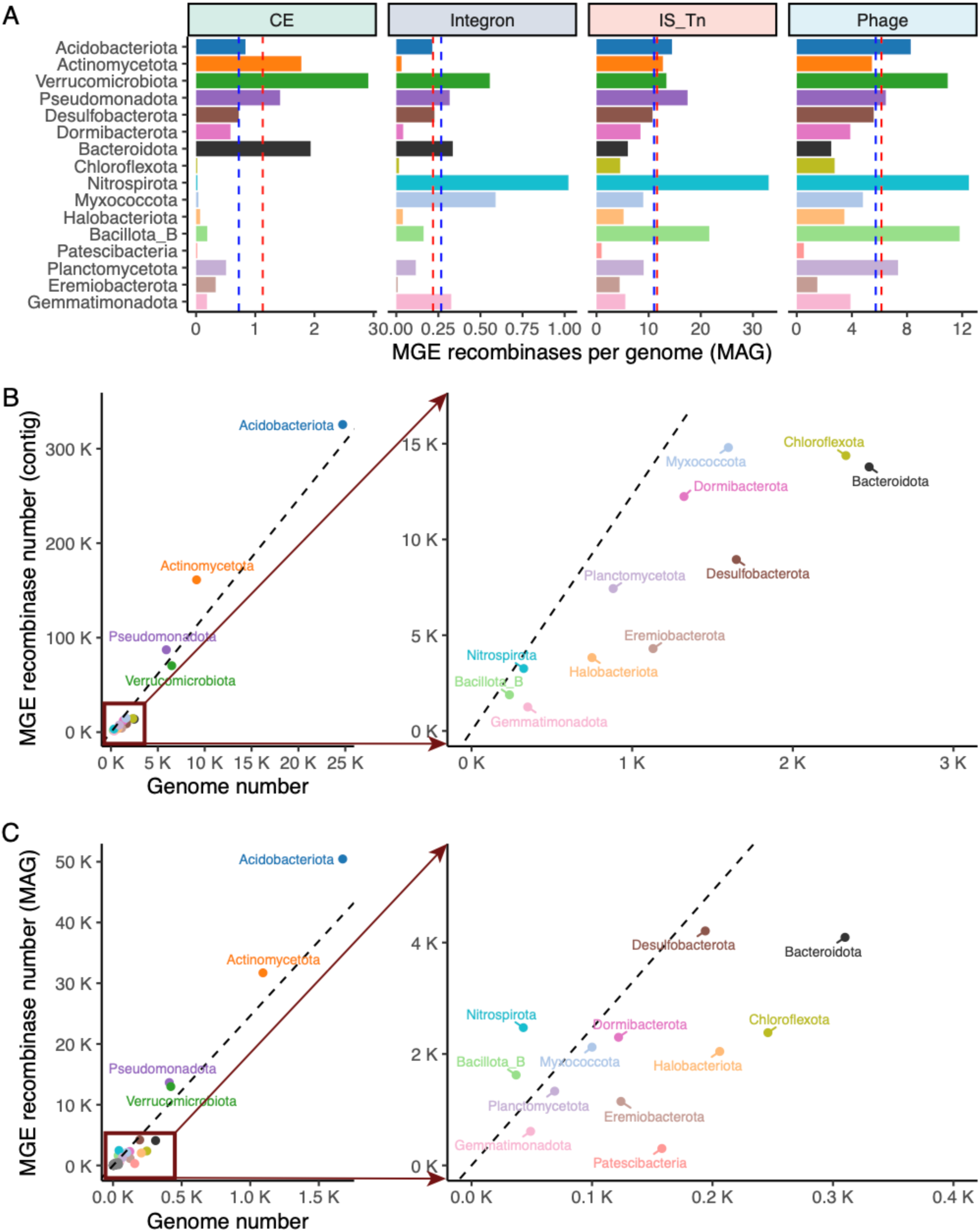
MGE recombinase and genome number distribution across phyla. **A**. MGE recombinase number per genome per phylum from MAGs. The dashed red lines are the average MGE recombinase number per genome across all phyla, and the blue dashed lines are the average across the phyla shown in the plot. **B.** Number of MGE recombinases against number of genomes per phyla using contigs. Dotted line represents the global average number of MGE recombinases per genome. The genome number is estimated from ribosomal genes.The right panel is a zoom-in of the area with cluttered dots in the left panel. **C.** Number of MGE recombinases against number of genomes per phyla using MAGs. The right panel is a zoom-in of the area with clustered dots in the left panel.

**Extended Data figure 6.**
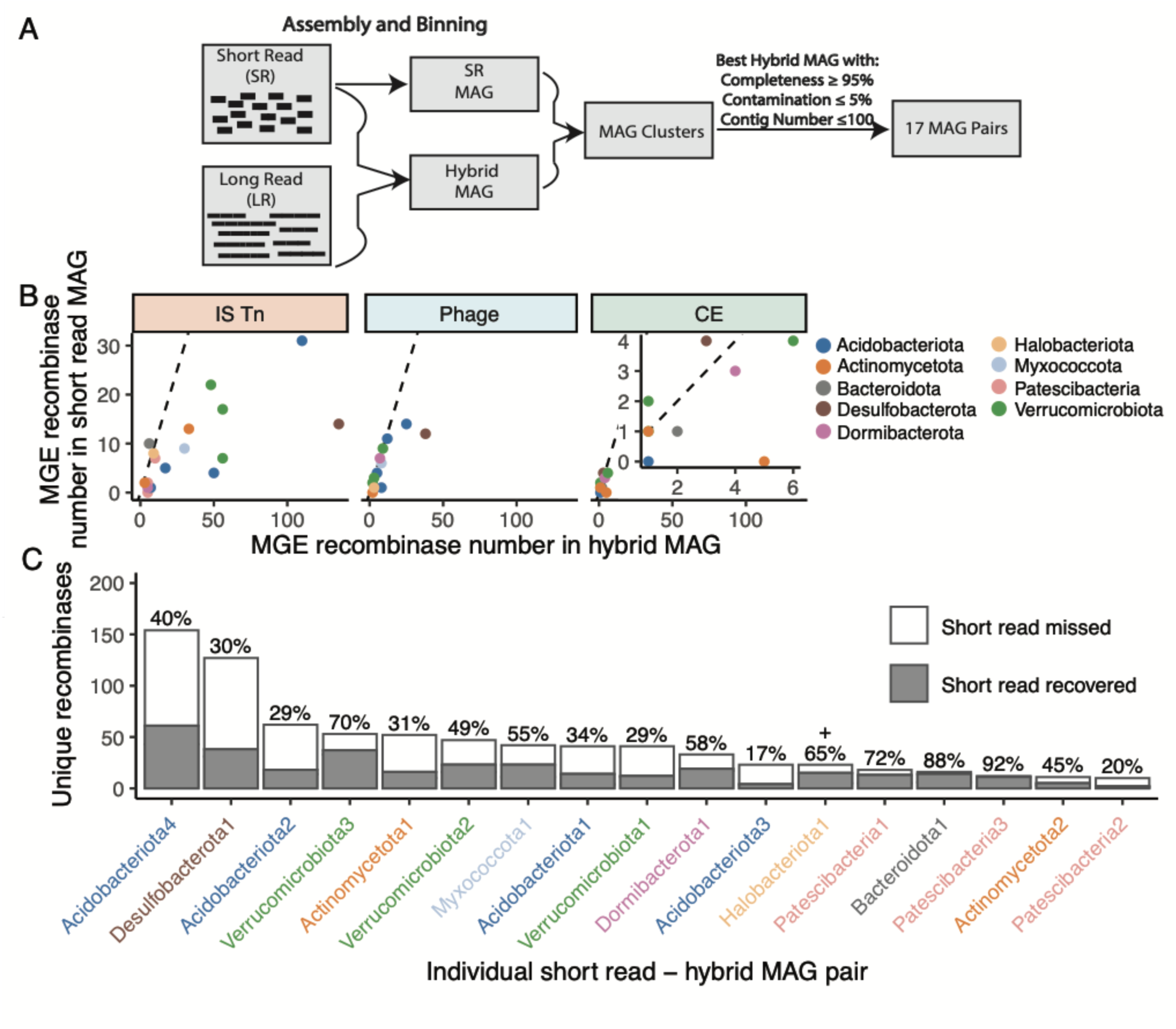
A six sample assessment of short-read assembly and binning artifacts for recovering MGE recombinases. **A.** Because short-read assemblies can struggle with hypervariable and/or repeat regions of genomes, six metagenomic samples were re-sequenced using high-fidelity long-read sequencing to develop a set of high-quality MAGs (>= 95% completeness, <=5% contamination, and <100 contigs) that could be compared for recombinase signal to short-read assembled MAGs from the same species cluster (>=95% ANI). A total of 17 such pairs resulted and were compared. **B.** Recombinase-centric patterns in recovery, where each circle represents the total recombinase number identified in the short read versus matching hybrid MAG and dashed lines are the 1:1 lines. The inset in CE is zoomin of clustered dots at lower left. **C.** Recombinase-centric patterns in recovery, where MGE recombinases are considered either recovered by short read or missed (i.e., if they were only found in hybrid MAGs) based upon searching via 100% identical protein sequence matches. **D.** Lineage-centric patterns in MGE recombinase recovery. MAG name color matches color in panel A. The numbers to the right of the bars are the percentages of total unique MGE recombinases recovered by short-read MAGs relative to the total.

**Extended Data figure 7.**
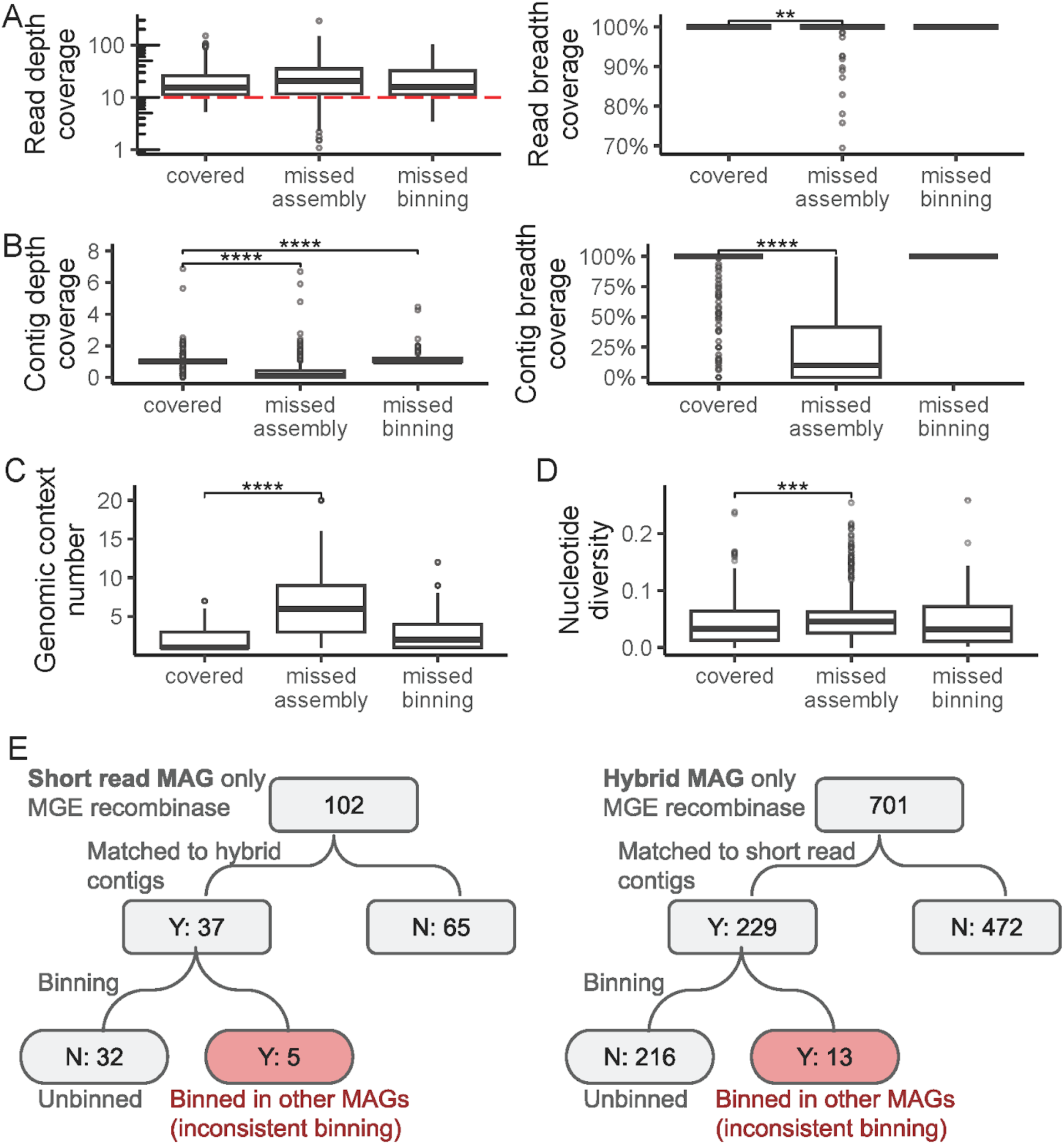
Technical assessment for why short-read MAGs miss some MGE recombinases. In all panels, “covered” represents MGE recombinases assembled and binned both in short-read and hybrid MAGs, “missed assembly” represents those in hybrid MAGs missing in short-read MAGs and short-read contigs, “missed binning” represents those in hybrid MAGs missing in short-read MAGs, but captured in short-read contigs (statistical significance per Fig.1). **A.** Short reads mapped to the full hybrid MAGs dataset of MGE recombinases evaluated by sequencing depth (left panel, vertical coverage) and breadth (right panel, horizontal coverage). The red dashed line at y=10 (left panel) approximates the minimal depth coverage needed for genome assembly considering the normal distribution of per bp position of depth coverages. **B.** Short-read-contig-based depth (left panel) and breadth (right panel) coverage per MGE recombinase to assess whether MGE recombinases were missed due to assembly or binning. **C.** To evaluate whether highly variable MGE-recombinase-caused genomic neighborhood context could prevent assembly, we quantified such variation as genomic context number via a clustering approach (see Methods). **D.** Nucleotide variation (pi, see Methods) across three MGE recombinase ‘capture scenarios’. **E.** Logic workflow to establish the fraction of short-(left panel) or hybrid-(right panel) MAG-only recombinases that are mis-binned in other MAGs. Each step is a yes or no answered question, and numbers represent MGE recombinase counts in our dataset.

**Extended Data figure 8.**
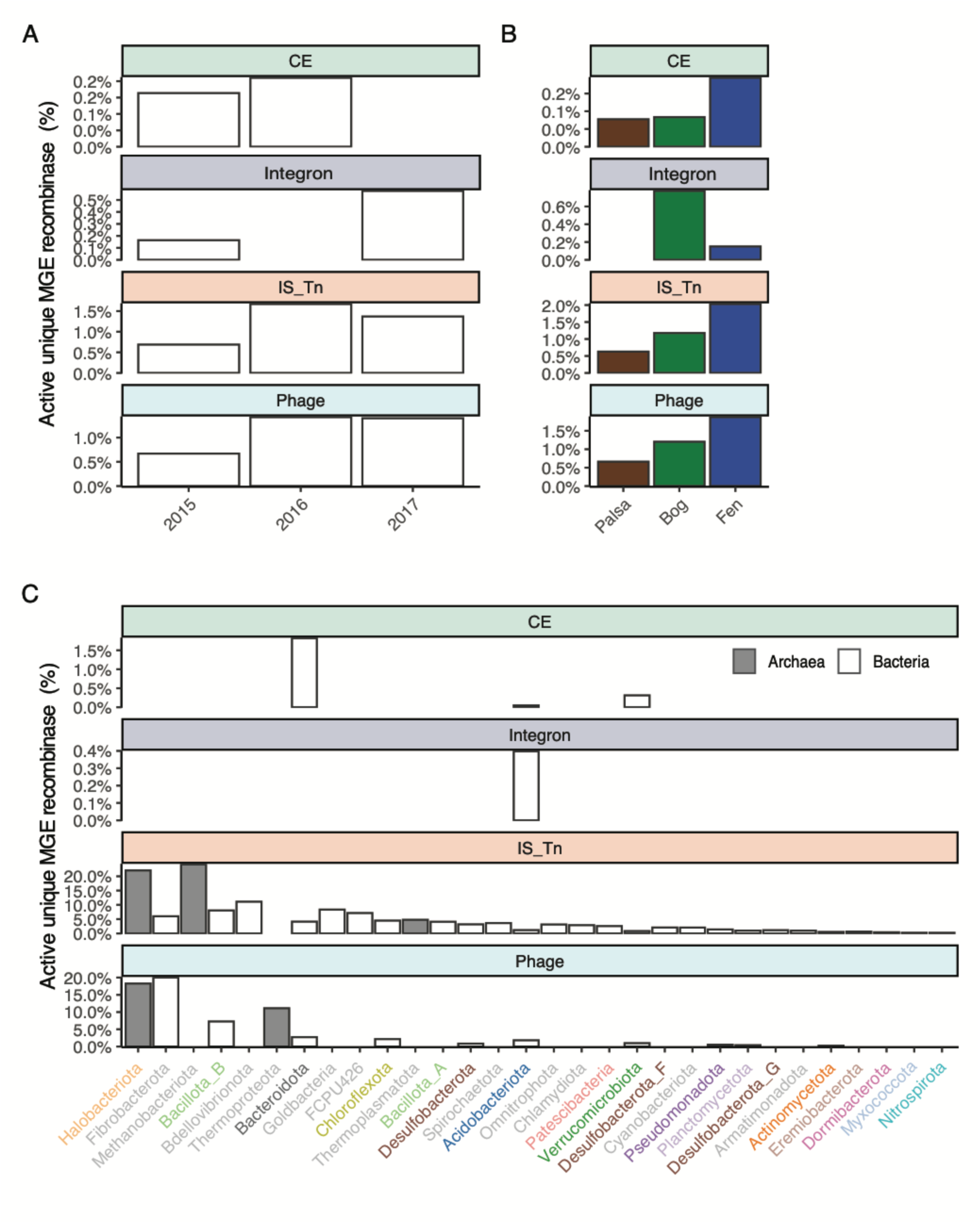
MGE recombinase activity accessed by metatranscriptomics. This figure complements Fig.5. **A.** Normalized active ratio among IS_Tn and Phage type recombinase types across host phyla. The rest of panels are MGE recombinase metatranscriptomics activity by active OTU (clustered at 100% identity in protein sequence) percentage. Each OTU with >=1 active MGE recombinase is considered as active. **B.** within different years from 2011 to 2017; **C.** within each habitat; **D.** within host phyla (only those >= 10 OTUs are shown; the X axis is ordered by the average of active OTU percentage across MGE recombinase types).

**Extended Data figure 9.**
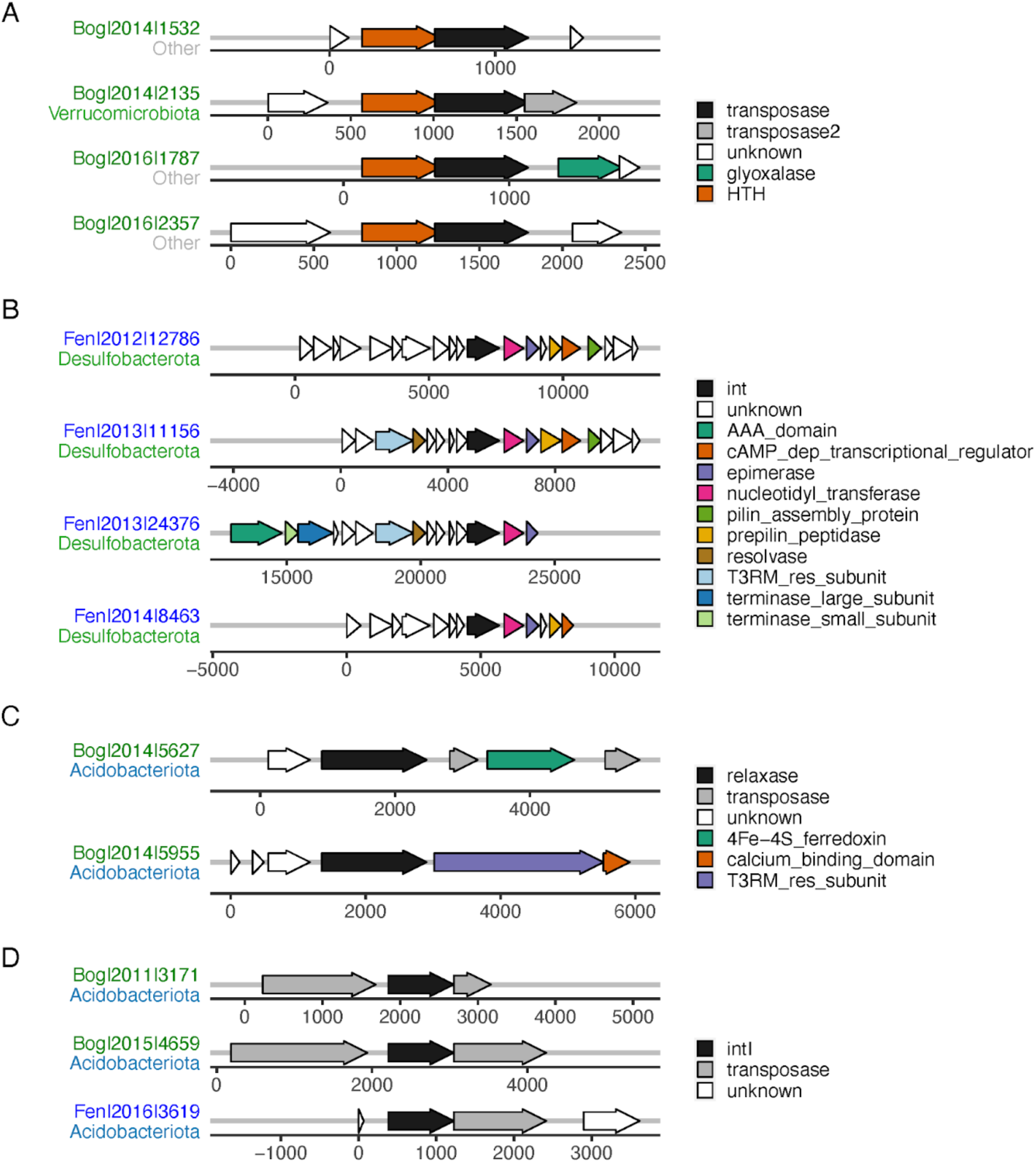
Example of unstable genomic neighborhoods of MGE recombinases. The labels at the left of genome maps are habitat, year, and contig length (separated by “|”), followed by host phyla. **A**. IS_Tn type (transposase); **B**. Phage type (int, short for integrase); **C**. CE type (relaxase), and **D**. Integron type (intI) recombinase.

**Extended Data figure 10.**
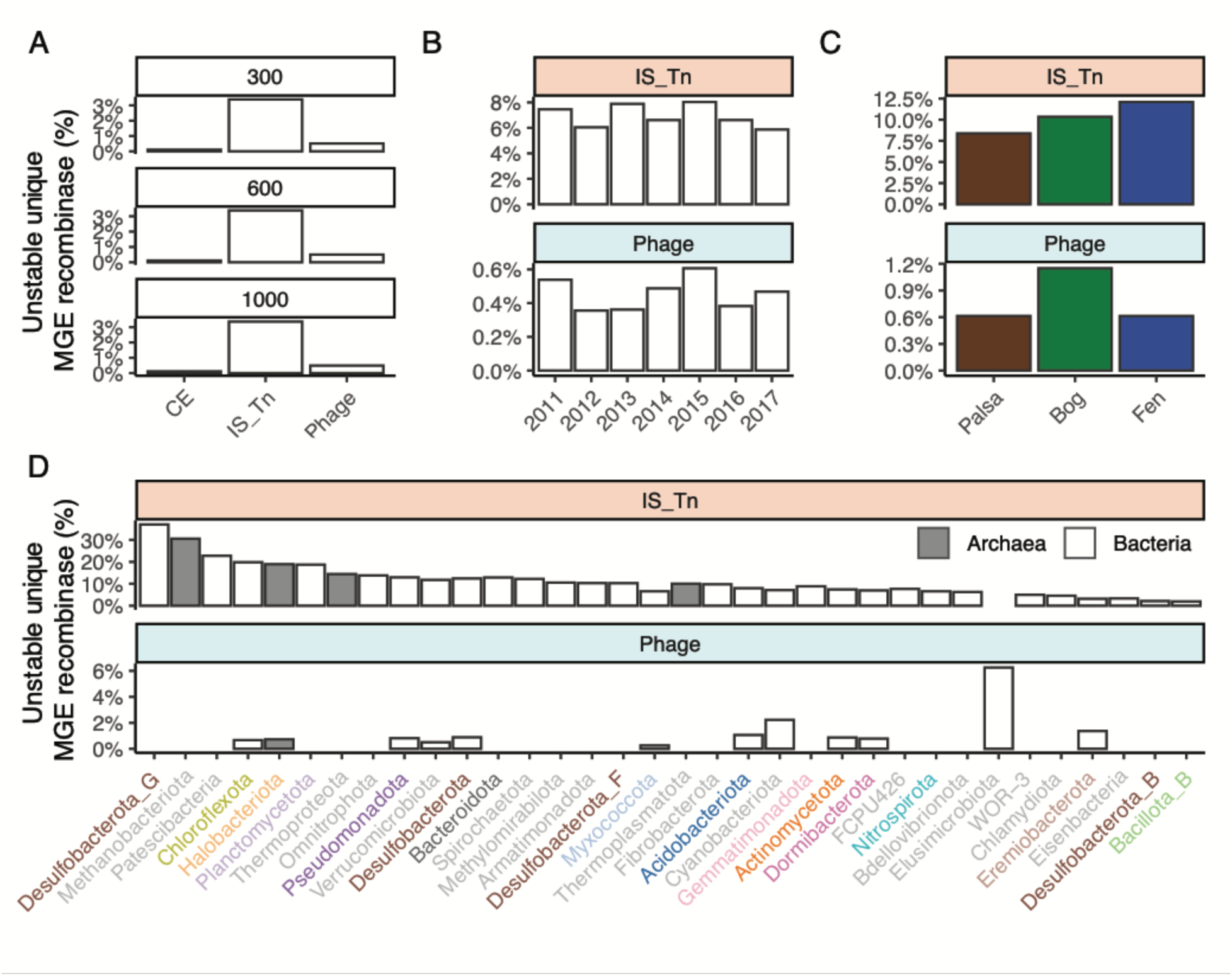
MGE recombinase genomic neighborhood stability. MGE activity could lead to their genomic neighborhood instability. To measure MGE recombinase genomic neighborhood stability, MGE recombinases are clustered at 100% AAI, and neighborhoods of 300 bp up/down-stream of recombinase are extracted. If different neighborhoods (<1% ANI) are detected within a cluster, the recombinase cluster is considered as unstable. The proportion of unstable MGE recombinase OTU were summarized: **A.** within each MGE type across neighborhood length cutoff ranging from 300 - 1000bp; **B.** within each year; **C.** within each habitat; and **D.** within host phyla (only those w/ >=10 OTUs are shown; the X axis is ordered by the average across MGE recombinase types). “CE” and “Integron” are not shown here due to lack of data.

**Extended Data figure 11.**
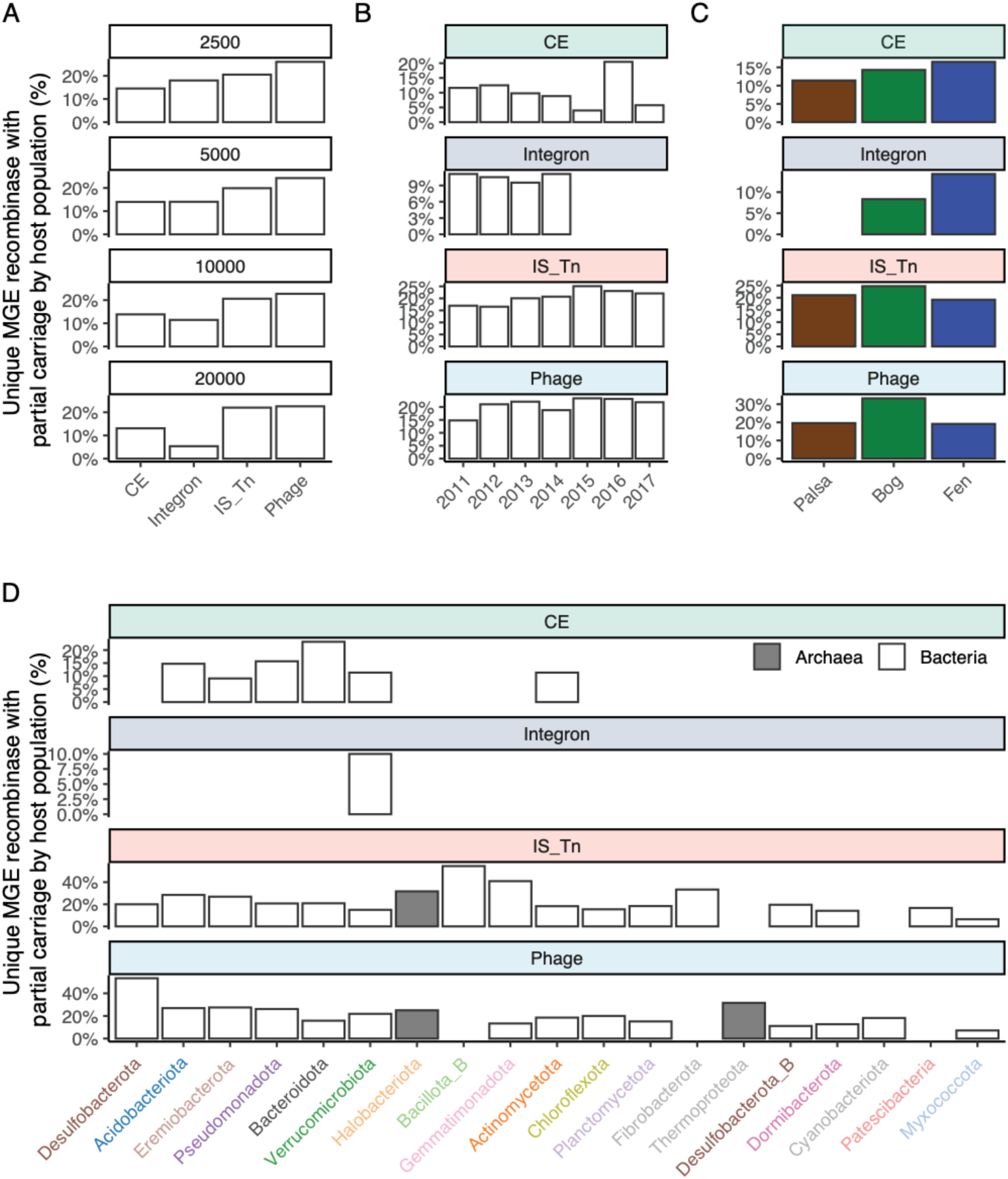
MGE recombinase partial carriage by microbial populations. MGE activity could lead to partial carriage of MGE in microbial populations. Here MGE recombinase partial carriage is defined as significantly lower coverage of an MGE recombinase than of its contig (only large contigs >=20kb upstream and downstream of a MGE recombinase are used to reduced the spurious coverage estimates from short contigs), and further MGE recombinase OTUs (100% AAI) with >= 1 partially carried MGE recombinase are considered as partially carried OTUs. The percentage of partially carried MGE recombinase OTUs were measured: **A.** within each MGE type across neighborhood length cutoff ranging from 2500 - 20,000bp; **B.** within each year; **C.** within each habitat; and **D.** within host phyla (only those w/ >= 10 OTUs are shown; the X axis is ordered by the average across MGE recombinase types).

