## Supplemental Text for "Mobile genetic elements that shape microbial diversity and functions in thawing permafrost soils"

### Supplementary Text

***Short reads under-detect active MGE recombinases, especially insertion sequences.***

As a technical control for sequencing artifacts, we sought to investigate the impact of short read assembly on MGE recombinase recovery from metagenomes as compared to that recovered from short- plus long-read hybrid assemblies. To evaluate this, we leveraged paired short-read and high-fidelity long-read sequencing data from 6 permafrost thaw soil samples collected in 2019 from Stordalen Mire. In this analysis, we sought to use high quality hybrid MAGs as the best available references to understand the impact of short read assembly and binning on MGE recombinase recovery.

We identified 17 high-quality hybrid-assembled MAGs (≥95% completeness, ≤5% contamination, <100 contigs) that also had representation in the short-read assemblies (clustered at 95% ANI). As expected, we found fewer MGE recombinases in short-read MAGs than in hybrid-assembled MAGs (n = 312 versus 937, respectively, across all MGE types). While MGE recombinase type recovery varied, IS_Tn type was consistently the least recovered (**Extended Data fig. 6B**). When directly comparing the recombinases detected in each pair of short-read and hybrid-assembled MAGs (100% protein identity clustering), the short-read MAGs recovered 57% CE, 35% IS_Tn, and 60% Phage recombinases in the 17 selected hybrid MAGs (**Fig. 2F**) (Integron recombinases were not included because of their low number, 27 in total in the shared MAGs), for an overall average of 36% (17–92% across individual MAGs, all MGE types combined) per hybrid MAG (**Extended Data fig. 6C**). Therefore, MGE recombinase recovery rate varies and is highly dependent on host taxon, but IS_Tn recombinases are the main type of recombinases under-detected in short-read datasets.

We next investigated why some MGE recombinases were recovered by hybrid-assembled MAGs, but missed by short-read MAGs. This revealed that virtually all hybrid-MAG MGE recombinases were found in short reads, but not short-read assembled contigs, suggesting that short read sequencing depth was not an issue, but assembly challenges prevented hybrid-assembled MGE recombinases from being captured in the short-read MAGs (**Extended Data fig. 7A,B**). We posited that assembly could fail either due to hypervariability of the recombinase locus itself and/or the surrounding genomic context. We found that recombinases missed by short read assembly ("missed assembly") have significantly higher numbers of genomic neighborhoods ($p \leq 0.0001$) and nucleotide diversity ($p \leq 0.001$, Wilcoxon rank-sum test) compared to those assembled and/or binned (**Extended Data fig. 7C,D**), which we interpret as confirming that hypervariable regions "break" short read assemblies due to mixed variation at that locus. For MGE recombinases successfully assembled from short reads, we found a relatively low number of inconsistent binning (<5%, **Extended Data fig. 7E**). This is consistent with previous studies (*20*, *21*) showing short read assembly and, to a lesser extent, binning under-detects MGE recombinases generally with simulated data. With the majority of the metagenomic data generated with short read sequencing, this means that the estimation of MGE prevalence and activity/mobility should be seen as a lower bound.

***Conservative MGE detection and impacted gene curation.***

To understand the capacity of the mobilome to affect microbial function across Stordalen, we sought to identify the genes whose distribution is influenced by mobilome activity, a group defined by both the cargo genes carried by an MGE and the host gene context into which the MGE is inserted. Because calling the precise boundaries of MGEs remains particularly challenging in metagenomic assemblies, we opted for a conservative approach, treating the outermost MGE hallmark genes as element boundaries and the genes between these bounds as cargo (Methods). Although this approach undercounts cargo genes, it can be seen as a lower bound. In our analyses – across integrons, CEs, and PhLs – we identified 90,219 cargo genes in all, annotated with 4,322 different KEGG functions (**table S3** item C,D,E,F). We took a similarly conservative approach to counting IS_Tn-impacted host genes, counting only those examples where the genomic neighborhood immediately upstream and downstream of IS/transposons could be robustly identified as the N- and C-termini of the same gene. Again, this approach will undercount IS_Tn-impacted host genes, as for instance insertion just upstream of a gene may influence or even block transcriptional regulation of that gene. Overall, we identified 12,332 interrupted host genes, annotated with 866 different KEGG functions. Then we selected and manually curated a subset (1,220) of the KEGG functions of specific interest to the Stordalen Mire thawing permafrost system for display based on functional categories, specifically Transport, Carbon/Nitrogen/Sulfur cycling, Signaling, Antiviral defense, Genome maintenance and expression, and prioritizing functions for which MGE impact was detected in multiple samples rather than single observations (**Fig. 4**, **table S4**).

***EMERGE field teams 2010-2017, 2019 and affiliations***

Darya Anderson[1], Kathryn Bennett[2], Sky Dominguez[1], Maria Florencia Fahnestock[2], Moira Hough[3], Joachim Jansen[4], Rhiannon Mondav[5], Apryl Perry[6], Gareth Trubl[7]

[1]Department of Environmental Science; University of Arizona; Tucson, AZ, USA.
[2]Department of Earth Sciences, University of New Hampshire; Durham NH, USA.
[3]Ecology & Evolutionary Biology, University of Arizona; Tucson, AZ, USA.
[4]Department of Geological Sciences, Stockholm University; Stockholm, Sweden.
[5]Australian Centre for Ecogenomics, School of Chemistry and Molecular Biosciences, University of Queensland; Brisbane, Queensland, Australia.
[6]Earth Systems Research Center, Institute for the Study of Earth, Oceans, and Space, University of New Hampshire; Durham, New Hampshire, USA.
[7]Department of Microbiology, The Ohio State University; Columbus, OH, USA.

***EMERGE coordinators and affiliations***

Patrick Crill[1], Jessica G. Ernakovich[2], Regis Ferriere[3], Mike Ibba[4], Malak M. Tfaily[5], Ruth K. Varner[6], Ahmed A. Zayed[7]

[1]Department of Geological Sciences and Bolin Centre for Climate Research, Stockholm University; Stockholm, Sweden.
[2]Department of Natural Resources and the Environment, University of New Hampshire; Durham, NH, USA.
[3]Department of Ecology and Evolutionary Biology, University of Arizona; Tucson, AZ, USA.

[4]Schmid College of Science and Technology, Chapman University; Orange, CA, USA.
[5]Department of Environmental Science; University of Arizona; Tucson, AZ, USA,
[6]Department of Earth Sciences and Earth Systems Research Center, University of New Hampshire; Durham, NH, USA.
[7]Department of Microbiology, The Ohio State University; Columbus, OH, USA.
